# Wnt-mediated interactions of tumor-initiating cells with a macrophage niche drive skin tumor formation

**DOI:** 10.1101/2020.07.17.209338

**Authors:** Silvia Fontenete, Johan Christensen, Adriana Martinez-Silgado, Eduardo Zarzuela, Javier Muñoz, Diego Megias, Donatello Castellana, Robert Loewe, Mirna Perez-Moreno

## Abstract

Skin tumor-initiating stem cells (tSCs) fuel skin squamous cell carcinoma (SCC) formation and development in a complex tumor microenvironment, but a role for immune cells in the tSC niche governing the process of tumor formation has remained elusive. Here, we define the existence of a tSC-macrophage niche, expressing high levels of Wnt ligands. Using conditional mouse genetic models to abrogate the secretion of Wnts, we show that both hair follicle SC- and macrophage-derived Wnts are essential for driving skin tumorigenesis, tSCs maintenance, and counteracting tumor regression. Loss of Wnts in either population uncouples the tSC-macrophage association. The proteomic signature of SCC cells reveals CD99 as a Wnt-dependent receptor for the macrophagecancer cell interaction. These results establish a role for a macrophage-tSC niche in governing SCC initiation and maintenance, uncovering potential candidates for immunoprevention against SCC.

**One Sentence Summary:** A macrophage-skin tumor-initiating cells bonding Wnt loop drives tumorigenesis.

## Introduction

Skin is subjected to constant exposure to mutagenic assaults, leading to the uncontrolled growth of tumor-initiated cells (tSC) supported by molecular signaling circuits^1^. Among them, Wnt/β-catenin signaling^2^ features prominently in the initiation and maintenance of tSCs^3,4^, while other signaling pathways drive SCC progression to invasiveness and malignancy (e.g., TGF-β)^5–7^.

The tumor immune microenvironment has well-established roles in governing the progression of initiated cells to SCC and conversion to malignancy^8–10^. Intriguingly, at initial stages, a correction of the abnormal growth of tSC leading to their regression has been observed in the absence of tumor promoters inducing inflammation^4,11^. However, whether the immune milieu in which tSCs reside sustains the abnormal growth of tSC leading to the initiation of SCC formation is not known.

Monocyte infiltration and skin macrophages activation form part of the earliest events in skin inflammation that lead to carcinogenesis^12^. Tumor-associated macrophages (TAMs) inhibit SCC antitumor responses^13^, are major sources of Wnt ligands in several tumors^14–17^, and recently TAMs expressing TGF have been associated in the transition from SCC *in situ* to invasive SCC^18^. However, whether macrophages at the tSC niche control the initial stages of tSCs proliferation and maintenance and exert this function as a source of Wnts is not known.

In this study, using SCC samples, mouse genetic models and proteomic analyses, we identify the existence of a TAM-tSC niche governed by a Wnt signaling loop. Both sources are required for tumor formation and to prevent tumor regression through the maintenance of tSC, CD99 being a receptor for the TAM-tSC interaction. Overall, these data establish that tSC Wnts drive the attraction of macrophages to the tumor niche, which in turn, as a source of Wnts, foster the initiation of skin tumors and sustain SCC stemness, opening new potential avenues for therapeutic approaches.

## Results

### tSC drive the attraction of TAMs to their niche in a Wnt-dependent manner

Tumor-associated macrophages (TAMs) in skin squamous cell carcinoma (SCC) foster tumor development and orchestrate anti-tumor immunity^12,13,15,19–23^. To assess the regional distribution of TAMs in the proximity of tSC, we conducted immunofluorescence microscopy analyses in mouse and human SCC, using the tSC master regulator Sox2^24,25^ as tSC marker, and the macrophage markers F4/80 and CD68^26^ (Fig. 1A). A high percentage of F4/80^+^ and CD68^+^ TAMs were found closely distributed in the vicinity of both mouse and human tSCs (<30 μm), 40 and 60%, respectively (Fig. 1B), uncovering the existence of a TAM – tSC niche in SCCs.

**Figure 1.**
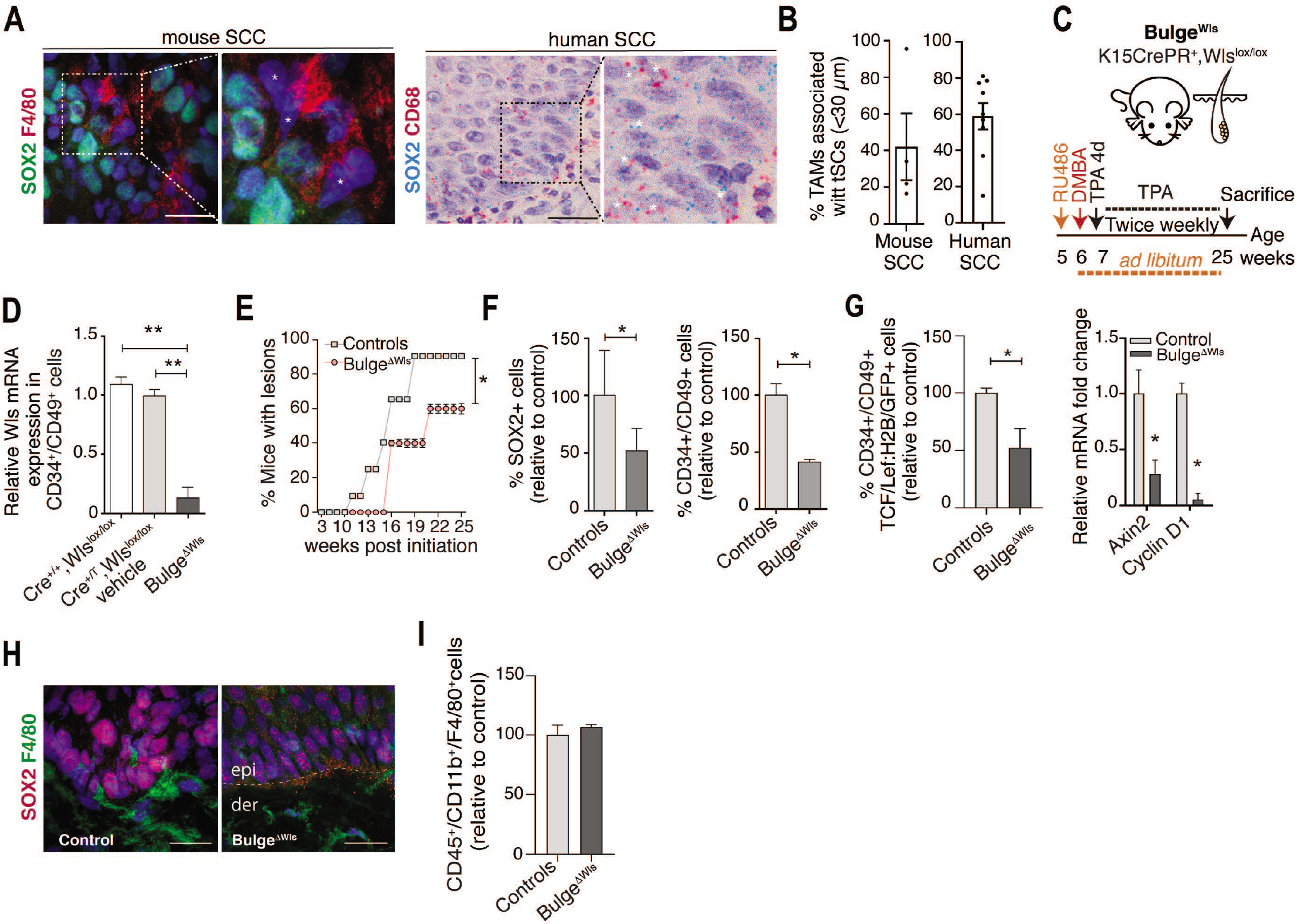
tSC drive the attraction of TAMs to their niche in a Wnt dependent manner. **(A)** Immunostaining for SOX2 (tumor initiating cells, tSCs, green) and F4/80+ macrophages (red) in mSCC *in situ* (Left). DAPI nuclear staining is represented in blue. Scale bar 20 μm. RNA *in situ* analysis for SOX2 (tSCs, blue) and CD68 (tumor associated macrophages, TAMs, red) in human SCC (hSCC) (Right). Scale bar 20 μm. Insets are magnified views. Asterisks denote positive stained TAMs. **(B)** Percentage of TAMs associated with tSCs (<30 μm) in mSCC (n=4) and hSCC (n = 8). **(C)** Protocol of Mifeprestone (RU486) and DMBA/TPA administration in the inducible mouse model K15CrePR1^+/T^ Wls^lox/lox^ (Bulge^Wls^). RU486 was injected before DMBA/TPA administration and maintained at *ad libitum* until the end of the experiments. Controls included mice carrying the Wls floxed alleles (Cre^+/+^, Wls^lox/lox^) and injected with RU486 and mice carrying all the alleles (Cre^+/T^; Wls^lox/lox^) and injected with vehicle. See Methods for more details. **(D)** qRT–PCR analysis of Wls mRNA expression in FACS-isolated CD34^+^CD49f^+^ tSC cells from the back skin of Bulge^Wls^ (n = 3) and control (n = 4) mice. **(E)** Quantification of mice with lesions after tumor initiation over the specified weeks in Bulge^ΔWls^ (n = 17) mice and control (n = 34) mice. **(F)** Quantification of SOX2^+^ immunostained tSCs numbers by IF in skin lesions in Bulge^ΔWls^ (n = 6) mice and control (n = 12) mice (Left). FACS quantification of CD34^+^CD49f^+^ tSCs in Bulge^ΔWls^ (n = 6) mice and controls (n = 7) mice (Right). **(G)** FACS quantification of CD34^+^CD49f^+^TCF/Lef:H2B/GFP^+^ tSCs in Bulge^ΔWls^ (n = 6) and control (n = 8) mice (Left). qRT–PCR analysis of the Wnt/β-catenin targets Axin2 and CyclinD1 expression in total skin of Bulge^ΔWls^ (n = 4) mice and control (n= 4) mice (Right) **(H)** Immunostaining for SOX2 (red) and F4/80 (green) in controls and Bulge^DWls^ mouse skin at the last point of the DMBA/TPA treatment. DAPI nuclear staining is represented in blue. Scale bar 20 μm. epi, epidermis; der, dermis. **(I)** FACS quantification of the proportion of CD45^+^CD11b^+^F4/80^+^ macrophages in Bulge^Wls^ (n = 4) and controls (n = 4) mice. Analyses were performed at the last time point of tumor development. **(D)** and **(G)** One-way Anova. **(E),(F)**,**(G)** and **(I)** Unpaired two-tailed Student’s *t*-test. All error bars represent s.e.m. *P < 0.05; **P < 0.01.

Next, we determined if tSC drive the recruitment of TAMs to their niche, and if TAMs localization at the tSC niche was associated with the formation and maintenance of initiated tumors. To this end, we targeted Wnt ligands, since Wnt signaling features prominently in SCC tumor initiation^2^, but the secretion of Wnt ligands in a non-inflamed environment eventually leads to tumor regression^4,11^. We used genetically engineered mouse models (GEMMs), and the well-established two-stage 7,12-dimethylbenz(a)anthracene (DMBA)/12-O-tetradecanoyl-phorbol-13-acetate (TPA) chemical skin carcinogenesis protocol^27,28^. TPA promotes the abnormal growth of genetic deregulated tSCs and inflammation^29^. The use of GEMMs allowed the abolishment of mature Wnts synthesis in specific populations, by breeding mice carrying floxed alleles of *Wntless (Wls)^30,31^* with mice carrying conditional promoters expressing Cre. We targeted *Wls* in HFSC using an inducible K15 promoter (K15CrePR1^+/T^ Wls^lox/lox^; Bulge^DWls^), since it has been demonstrated that upon DMBA/TPA oncogenic mutations, this population is able to initiate skin tumorigenesis^1,32,33^ (Fig. 1C). RT-PCR analyses of FACS-isolated cells validated the reduction of Wls mRNA expression in bulge CD34^+^CD49^+^ cells (Fig. 1D). To provide evidence of a causal role of Wnt secretion by tSC in tumor initiation we ablated *Wls* before starting the DMBA/TPA skin carcinogenesis protocol (Fig. 1C). We observed a significant reduction in the percentage of mice with lesions (Fig. 1E and Fig. S1A) and histopathological characteristics (Fig. S1B). These results were associated with a decrease in the epithelial Sox2^+^ and CD34^+^ CD49^+^ tSC pools (Fig. 1F and Fig. S2C). The reduction in SOX2^+^ tSCs numbers was accompanied by a decrease in SOX2^+^ cells proliferation (Fig. S1D, S1E) and promoted epidermal differentiation (Fig. S1F, S1G).

Although we cannot formally exclude that a combined involvement of canonical and non-canonical Wnt signaling in HFSC underlies this effect^34,35^, we explored the influence of the loss of Wnts secretion in the activation of Wnt/β-catenin signaling in tSC by breeding the GEMMs under the background of a TCF/Lef:H2B-GFP reporter mouse line and observed that CD34^+^/CD49^+^ TCF/Lef:H2B-GFP^+^ tSC decreased in number (Fig. 1G), as did the expression of Wnt/β-catenin downstream target genes in skin (Fig. 1G). Interestingly, these reductions were accompanied by a reduction of TAMs recruitment to tSC sites, where only 17% TAMs were associated to the tSC niche, without changes in total skin TAMs numbers (Fig 1H, I). Taken together, these data reveal that Wnts secreted by tSC are required for supporting the initiation of skin SCC, an event closely associated with the recruitment of TAMs to their microenvironment.

### Macrophage-Wnts are essential for the distribution of TAMs to the tSC niche and promote tumor formation

Mammalian Wnts act as short-range signals^36^, and TAMs can respond to and be a source of Wnt ligands in several tumors^19,34,37^. To assess whether TAMs that were distributed in the vicinity of tSC also express Wnt ligands, we conducted double mRNA in situ hybridization assays in a panel of human SCCs, using the macrophage marker CD68^26^. CD68^+^TAMs and cancer cells expressed several Wnt ligands throughout the skin (Fig. 2A-C and Fig. S2A-C). Notably, CD68^+^TAMs closely associated with cancer cells highly expressed Wnt7b and Wnt10a, previously linked to SCC as canonical Wnt ligands^38,39^, compared to Wnt3a (Fig. 2A, B). Cancer cells closely associated with TAMs also expressed Wnt3a, Wnt7b, and Wnt10a (Fig. 2A, C). Overall, these results expose that both tSC and TAMs express Wnt ligands at the tSC niche.

**Figure 2.**
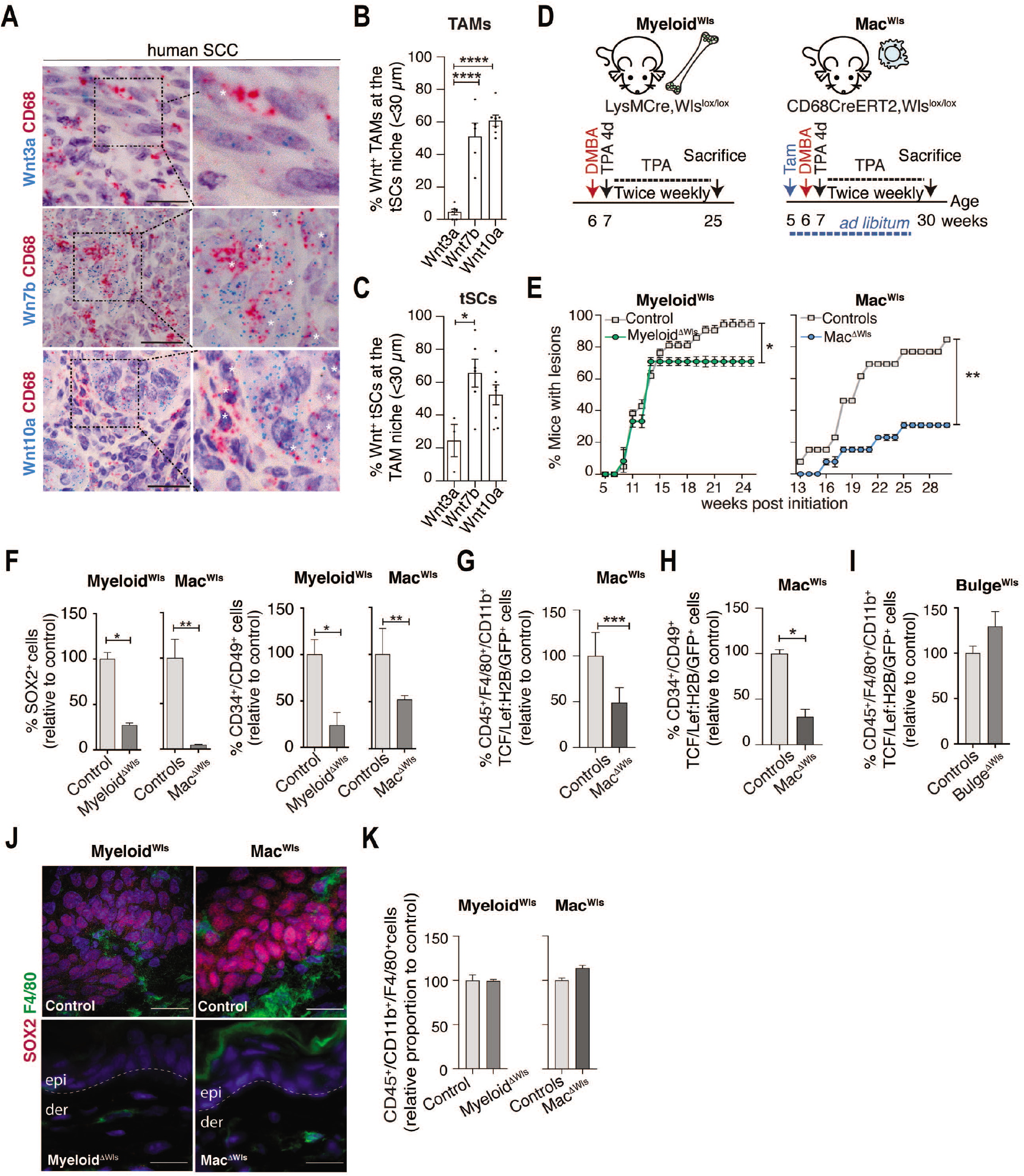
TAM-derived Wnts promote skin tumor initiation and development. **(A)** RNA *in situ* analysis for CD68 (TAMs, red) and Wnt3a (blue, top) Wnt7b (blue, middle) and Wnt10a (blue, bottom) in hSCC. Scale bar 20 μm. Insets are magnified views. Quantification of TAMs **(B)** and tSCs **(C)** in hSCC (n = 3-8) expressing Wnt3a, Wnt7b or Wnt10a mRNA (> 3+ dots) in the proximity (< 30 μm) of SOX2^+^tSC or CD68^+^TAMs, respectively. **(D)** Protocol of DMBA/TPA and Tamoxifen (TAM) administration used in the constitutive LyMCre^+/T^ Wls^lox/lox^ (Myeloid^Wls^) and the inducible CD68CreERT2^+/T^ Wls^lox/lox^ (Mac^Wls^) mice models. TAM was injected for 4 consecutive days in the Mac^Wls^ mice before DMBA/TPA administration and maintained at *ad libitum* until the end of the experiments. Controls included mice carrying the Wls floxed alleles (Cre^+/+^, Wls^lox/lox^) (Myeloid^Wls^) and injected with TAM (Mac^Wls^). For Mac^Wls^, additional controls included mice carrying all the alleles (Cre^+/T^; Wls^lox/lox^) and injected with vehicles. See Methods for more details. **(E)** Quantification of mice with lesions after tumor initiation over the specified weeks in Myeloid^ΔWls^ (n = 24) (Left) and Mac^ΔWls^ (n = 16) (Right) mice and control (n = 24, 11) mice, respectively. **(F)** Quantification of SOX2^+^ immunostained tSCs numbers by IF in skin lesions in Myeloid^ΔWls^ (n = 10) and Mac^ΔWls^ (n =6) mice and controls (n=4,6), respectively (Left). FACS quantification of CD34^+^CD49f^+^ tSCs in Myeloid^ΔWls^ (n = 12) and Mac^ΔWls^ (n =6) mice and controls (n=15,8), respectively (Right). **(G)** FACS quantification of CD45^+^CD11b^+^F4/80^+^TCF/Lef:H2B/GFP^+^ TAMs in Mac^ΔWls^ (n=12) and control (n=12) mice. **(H)** FACS quantification of CD34^+^CD49f^+^TCF/Lef:H2B/GFP^+^ tSCs in Mac^ΔWls^ (n = 12) and control (n = 12) mice. **(I)** FACS quantification of CD45^+^CD11b^+^F4/80^+^TCF/Lef:H2B/GFP^+^ TAMs in Bulge^ΔWls^ (n = 13) and control (n = 8) mice. **(J)** Immunostaining for SOX2+ (red) and F4/80+ (green) cells in control and Myeloid^ΔWls^ and Mac^ΔWls^ mouse skin at the last point of the DMBA/TPA treatment. DAPI nuclear staining is represented in blue. Scale bar 20 μm. **(K)** FACS quantification of total CD45^+^CD11b^+^F4/80^+^ macrophages isolated from the back skin of Myeloid^ΔWls^ (n = 12) and Mac^ΔWls^ (n = 8) mice and control (n= 15,13) mice respectively. Analyses were performed at the last time point of tumor development. **(B)** and **(C)** One-way Anova. **(E)**, **(F)** and **(K)** Unpaired two-tailed Student’s *t*-test. **(G)**, **(H)** and **(I)** Unpaired two-tailed Student’s *t*-test with Welsh’s correction. All error bars represent s.e.m. *P < 0.05; **P < 0.01; ****P < 0.0001.

To determine a causal role for the secretion of Wnt ligands by TAMs in tumor initiation, we turned again to mouse genetics and the two-stage DMBA/TPA chemical skin carcinogenesis protocol. We targeted *Wls* in myeloid cells (constitutive LysMCre^+/T^;Wls^lox/lox^; Myeloid^ΔWls^) and macrophages (inducible CD68CreERT2^+/T^;Wls^lox/lox^; Mac^ΔWls^). RT-PCR analyses of FACS-isolated cells validated the reduction of Wls mRNA expression in both cell populations (Fig. S4A). The ablation of *Wls* before conducting the DMBA/TPA skin carcinogenesis protocol led to a significant decrease in tumor formation (Fig. 2E and Fig. S3A), and almost a complete abrogation of histopathological characteristics (Fig. S3B). These effects were more pronounced than those observed in Bulge^ΔWls^ mouse skin (Fig. S1B). The prevention of tumor initiation in Myeloid^ΔWls^ and Mac^ΔWls^ skin was associated with an almost total decrease in the epithelial SOX2^+^ and CD34^+^ CD49^+^ tSC pools (Fig. 2F and Fig. S4B), accompanied by a reduction in SOX2^+^ cells proliferation (Fig. S4D and S4E) and an increase in epidermal differentiation (Fig. S4F and S4G). Interestingly, Wnt/β-catenin signaling was not only reduced in F4/80^+^ cells (Fig. 2G), but also in the epithelial CD34^+^CD49^+^ tSC pool (Fig. 2H), and in the total skin of Myeloid^ΔWls^ and Mac^ΔWls^ mice (Fig. S4C). However, the converse influence of tSC-Wnts over TAMs Wnt/β-catenin activity was not observed in Bulge^ΔWls^ mice (Fig. 2I). The almost complete absence of SOX2+ tSCs and reductions in Wnt signaling precluded the localization of F4/80+ cells to lesion sites, even when the TPA inflammatory stimulus was maintained (Fig. 2J), without changes in their total numbers (Fig 2K). Taken together, these data revealed three important points regarding Wnts secreted by TAMs in mouse skin SCC: 1) they are required for supporting the initiation and maintenance of skin SCC, 2) they contribute to the activation of β-catenin signaling and proliferation of skin tSCs and 3) TAM-derived Wnts sustain β-catenin signaling in tSC, but not conversely.

### Loss of mature Wnts in either tSCs or TAMs leads to tumor relapse

To assess the potential therapeutic relevance of inhibiting the secretion of mature Wnts in tSCs or TAMs in their native skin tumor microenvironment by promoting tumor regression, we allowed the development of DMBA/TPA-induced skin tumors “in situ” before ablating *Wls* in Bulge^ΔWls^, Myeloid^ΔWls^, Mac^ΔWls^ or in tSCs/TAMs (Bulge^ΔWls^/Mac^ΔWls^) mice (Fig. 3A). The results showed that tumors regressed in all mouse models after 2 weeks of treatment (Fig. 3B and Fig. S5A), compared to their control counterparts. Mac^ΔWls^ skin exhibited a higher therapeutic effect in tumor regression than the one observed in Bulge^ΔWls^ skin, while the impact of the deletion of Wls in Bulge^ΔWls^/Mac^ΔWls^ induced a complete relapse of skin lesions, consistent with the histopathological analyses of mouse skin samples from each experimental set up (Fig. S5B). Overall, these results underscore the relevance of the secretion of Wnts from either tSCs or TAMs to sustain the maintenance of initiated skin tumors. In a similar scenario to the one observed during tumor initiation, the loss of *Wls* in Bulge^ΔWls^, Myeloid^ΔWls^, Mac^ΔWls^ or in Bulge^ΔWls^/Mac^ΔWls^ mice bearing tumors led to a decrease in the epithelial SOX2^+^ and CD34^+^ CD49^+^ tSC pools (Fig. 3C and Fig. S6A), reductions in SOX2^+^ cells proliferation (Fig. S6B and 6C), a decrease in the expression of Wnt/β-catenin downstream target genes (Fig. S6D), and promoted epidermal differentiation (Fig. S6E and S6F), without signs of apoptosis, except for Bulge^ΔWls^Mac^ΔWls^ mouse skin (Fig. S6G). Moreover, as observed during tumor initiation, the loss of Wnts secretion in tSC led to a decrease in Wnt/β-catenin signaling in tSC but not in TAMs (Fig. 3D, E), while TAMs Wnts were required for the activation of β-catenin signaling in both TAMs and tSCs (Fig. 3D, E). The loss of Wnts in either cell source led to a complete uncoupling of their interactions at the tSC niche (Fig. 3F), despite sustaining a TPA tumor-promoting environment, without changes in total F4/80^+^ cell numbers (Fig. 3G). Overall, these results reveal the requirement for both tSC- and TAM-derived Wnts for the maintenance of skin SCC and indicate a role for TAM-derived Wnts in governing SCC stemness.

**Figure 3.**
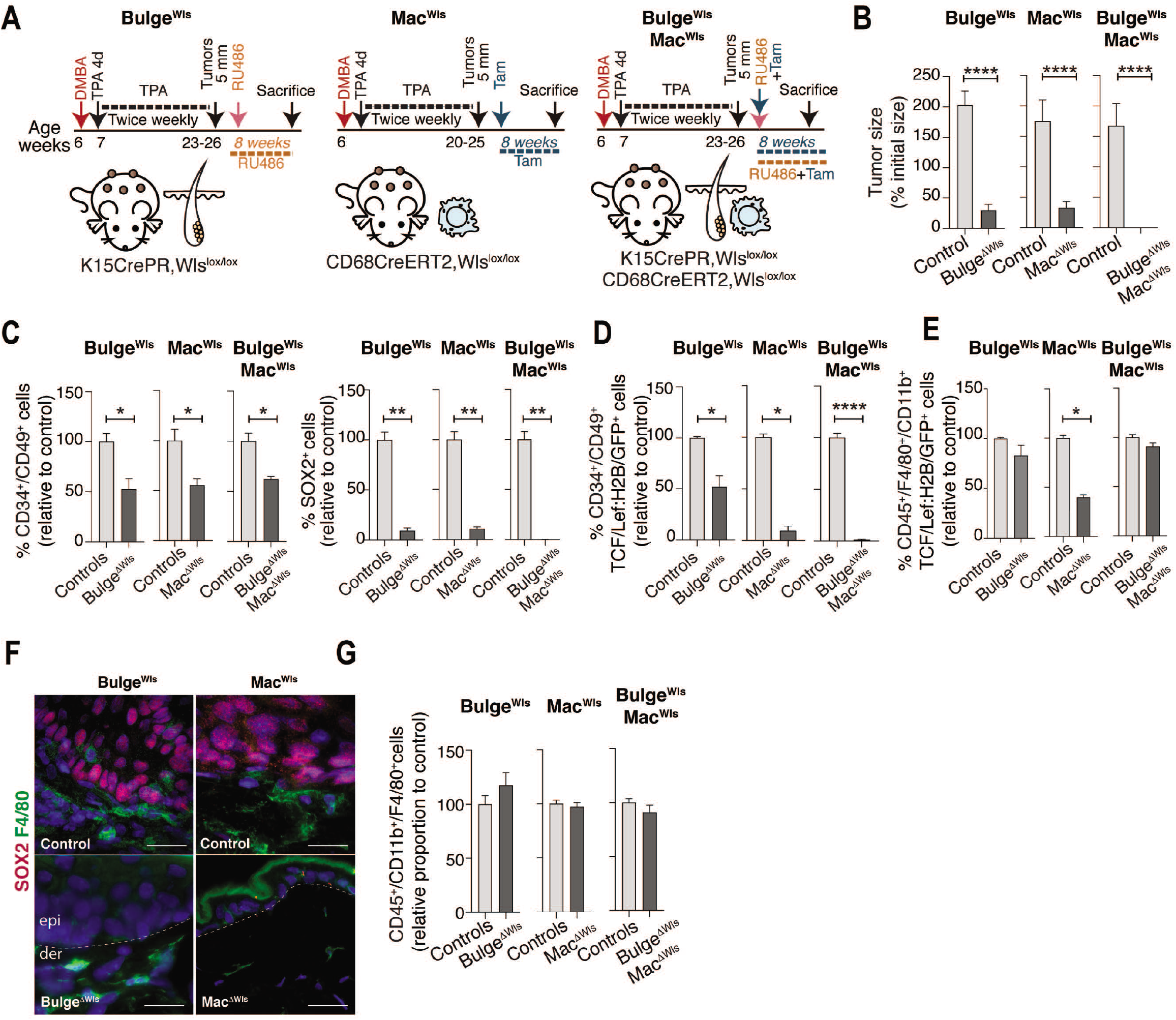
Loss of Wnt from tSCs and macrophages leads to tumor regression. **(A)** Protocol of Tamoxifen or/and RU486 administration in mice with DMBA/TPA-induced skin lesions. See Methods for more details. **(B)** Tumor size 8 weeks after RU467, TAM or RU486+TAM administration in Bulge^ΔWls^ (n = 9), Mac^ΔWls^ (n = 9), and Bulge^ΔWls^Mac^ΔWls^ (n = 3) mice and controls (n = 12, 12, 4), respectively. **(C)** FACS quantification of CD34^+^CD49f^+^ tSCs in Bulge^ΔWls^ (n = 6), Mac^ΔWls^ (n = 6) and Bulge^ΔWls^Mac^ΔWls^ (n = 4) mice and control (n = 8,7,3) mice, respectively (Left). Quantification of SOX2^+^ immunostained tSC numbers by IF in skin lesions (n = 3 mice in each group) (Right). **(D)** FACS quantification of CD34^+^CD49f^+^TCF/Lef:H2B/GFP^+^ tSCs in Bulge^ΔWls^ (n = 6), Mac^ΔWls^ (n = 6) and Bulge Mac^ΔWls^ (n = 3) mice and control (n = 8, 8, 8) mice, respectively. **(E)** FACS quantification of CD45^+^CD11b^+^F4/80^+^TCF/Lef:H2B/GFP^+^ TAMs in Bulge^ΔWls^ (n = 13), Mac^ΔWls^ (n = 6) and Bulge^ΔWls^ Mac^ΔWls^ (n = 3) mice and control (n = 8, 8, 8) mice, respectively. (**F**) Immunostaining for SOX2^+^ and F4/80^+^ cells in Bulge^DWls^ and Mac^DWls^ mouse skin at the last point of tumor regression. DAPI nuclear staining is represented in blue. Scale bar 20 μm. **(B)** Mann-Whitney test. **(C), (D)**, **(E)** and **(G)**. Unpaired two-tailed Student’s *t*-test. All error bars represent s.e.m. *P < 0.05; **P < 0.01; ****P < 0.0001.

### Direct Wnt-driven tSC-TAM interactions

Our prior results indicated that Wnts are required for the attraction of TAMs to the tSC niche. To address if a direct tSC – TAMs communication underlies the observed effect, we turned to *in vitro* studies (Fig. 4A) using the mSCC derived tumor cell line HaCa4 (TC)^40^ that expresses both CD34 and Sox2 tSC markers (Fig. S7A), and the F4/80^+^ CD11b^+^ mouse macrophage line RAW 267.4 (MØ)^41^ (Fig. S7B). Similar to the *in vivo* scenario, both TC and MØ expressed Wnts (Fig. S7C). Assessing the requirement for the secretion of Wnt ligands was made possible by pre-treating cells with the inhibitor IWP-2^42^ compared to the vehicle, which led to a reduction of the expression of Wnt ligands (Fig. S7D) and the expression of Wnt/β-catenin signaling target genes (Fig. S7D). In seeking understanding of whether the TC – MØ heterotypic interactions maintain tSC characteristics, we investigated if Wnt secreted by TC or MØ influenced the expression of the tSC markers Sox2 and CD34 using conditioned media (CM) (Fig. 4 A, B). Compared to the effect of TC CM, MØ CM significantly increased the expression of CD34 and SOX2 in TC that was reduced under IWP2 pretreatment conditions (Fig. 4B). These results were in agreement with the observed reductions in colony formation (Fig. 4C). We also investigated the influence of either cell population in stimulating the expression of Wnt ligands and the expression of Wnt/β-catenin targets on their opposite cell type when growing in co-culture. Both Wnt cellular sources were required to stimulate their reciprocal expression of Wnts (Fig. S7E). Moreover, MØ Wnts were necessary for the expression of β-catenin signaling targets in TC, but not conversely (Fig. S7F) in agreement with the *in vivo* findings (Fig. 2G-I).

**Figure 4.**
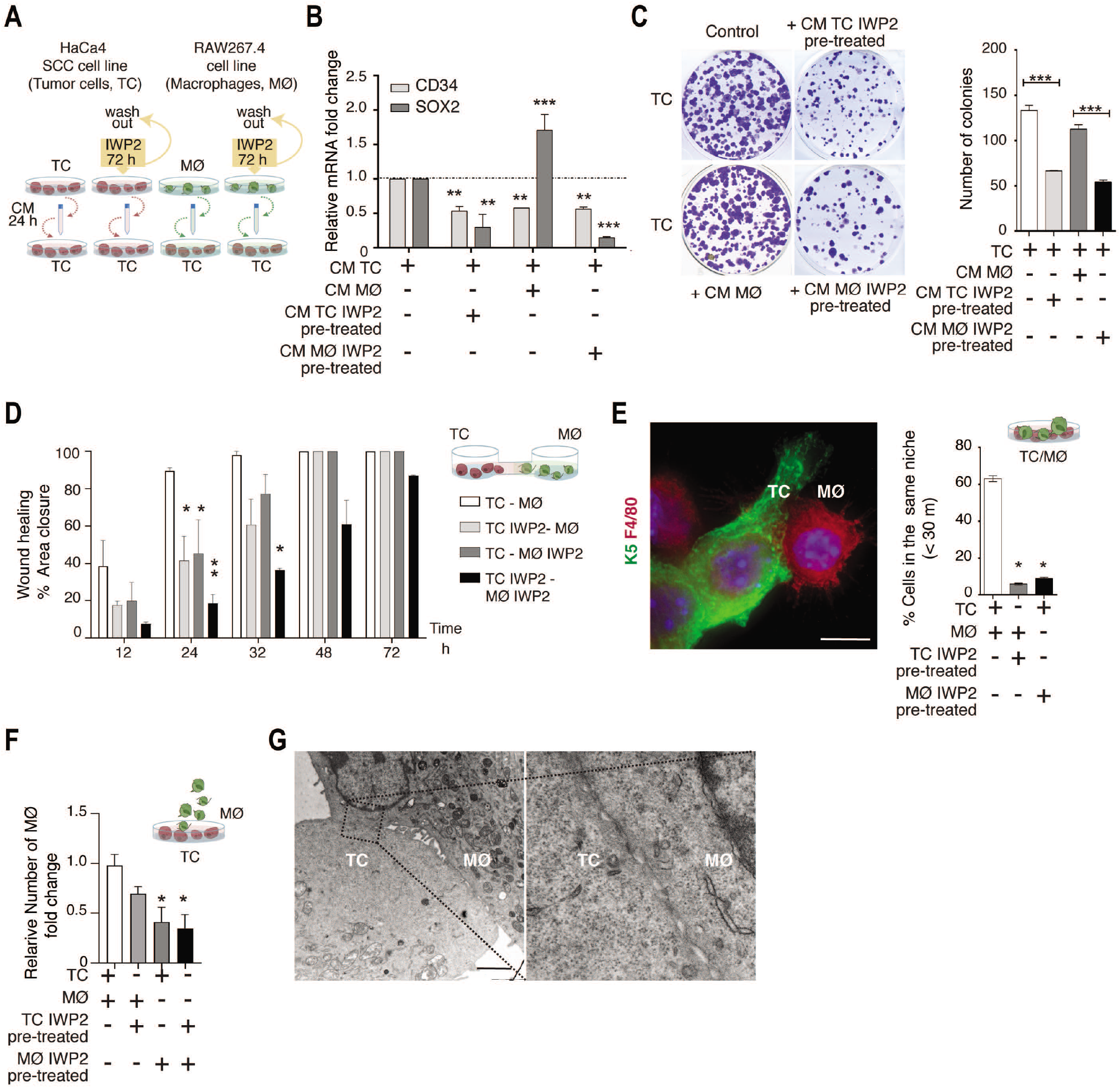
A Wnt signaling loop between cancer cells and macrophages sustains tSC features and promotes their attraction and direct association. **(A)** Protocol used to stimulate HaCa4 mSCC cells (Tumor cells, TC) with TC-or RAW264.7 cells (Macrophages, MØ) – conditioned media (CM) harvested from cells pre-treated with IWP-2 or vehicle. **(B)** qRT–PCR analysis of CD34 and SOX2 mRNA expression in TC stimulated with the different CM depicted. Data were normalized to CM TC values (n = 3) (bottom). **(C)** Colony formation assay of TC stimulated with CM from the different cellular sources depicted (Left). Quantification of the number of TC colonies (n = 3) (Right). **(D)** Wound area closure between TC and MØ under vehicle or IWP2 pre-treatment conditions, relative to the starting time point (n = 3). **(E)** Immunostaining for K5 (TC, green) and F4/80 (MØ, red) (Left). Scale bar 10 μm. Quantification of MØ in the TC niche (<30 μm) under vehicle or IWP2 pre-treatment conditions (n = 3) (Right). **(F)** FACS quantification of adhered MØ to TC under vehicle or IWP2 pre-treatment conditions (n = 3). **(G)** Transmission electron microscopy image of TC and MØ co-cultures. Scale bar 2 μm. Right panel represents a magnified inset. **(B), (C), (E) and (F)** One-way Anova, **(D)** Two-way Anova. n represents the number of independent experiments. All error bars represent s.e.m. *P<0.05; ** P<0.01, *** P<0.001, **** P<0.0001.

Live cell-imaging revealed that pre-treating either TC or MØ or both populations with IWP2 abrogated their attraction, decreasing their ability to migrate towards each other (Fig. 4D). Interestingly, under control conditions, TC and MØ dynamically collided forming heterotypic contacts to seal the gap, an effect prevented by the absence of mature Wnts (See Movie S1). We confirmed the existence of Wnt-dependent interactions between TC and MØ by immunofluorescence analyses that showed the presence of closely associated TC and MØ within cell colonies in co-culture (Fig. 4E). The IWP2 pre-treatment of each population precluded their close association (Fig. 4E) and their heterotypic adhesion (Fig. 4F and Fig. S7G). The observed interactions between TC and MØ were direct, as analyzed by transmission electro-microscopy (Fig. 4G). Overall, these results expose the causal requirement of Wnt ligands for mediating the cancer cell – macrophage attraction and interaction and a causal role for macrophages in sustaining tSC features and Wnt/β-catenin activity.

### tSC membrane protein signatures modulated by TAM-derived Wnts

To identify potential cell adhesion molecules mediating the Wnt-dependent TC-MØ heterotypic interaction, we conducted a quantitative mass-spectrometry-based proteomic profiling of membrane-associated proteins in TC stimulated with MØ CM. The CM was obtained from MØ pre-treated with either IWP2 or the vehicle as control (Fig. 5A). Gene Ontology analyses of the total 2446 identified proteins from all conditions validated a prominent enrichment of membrane and plasma membrane proteins, cell junction, cell-cell junction, cell substrate junction, and actin cytoskeleton cellular component categories (Fig. 5B). Among the 775 identified plasma membrane proteins, a similar proportion of proteins was found upregulated or downregulated by MØ Wnts, when comparing IWP2 versus vehicle conditions (FDR <0.05) (Fig. 5C). Gene set enrichment analyses further revealed a leaning decrease of the Cell Adhesion Molecules (CAMs) category (p-value < 0.059, FDR 0.53) (Fig. 5D). The identified plasma membrane candidates comprised intercellular adhesion proteins (e.g., CD99, Pvrl2, DSP, JUP, JAM), membrane receptors (e.g., CD14, CD147, CD109), and focal adhesion and matrix remodeling membrane proteins (e.g., Itgb3, Itgb5, Adam9, Adam 17, CD147, Plaur). RT-PCR analyses to quantify their differential expression in TC, when stimulated with MØ CM under IWP2 versus vehicle conditions, revealed a significant differential expression of CD109, a GPI protein involved in skin carcinogenesis^43^, and the cell-cell adhesion molecules Pvrl2^44^ and CD99^45,46^ whose roles in SCC are not understood (Fig. 5E). These results support a role for MØ Wnts in inducing the expression of cell membrane proteins in TC potentially involved in mediating the adhesive interactions between cancer cells and TAMs.

**Figure 5.**
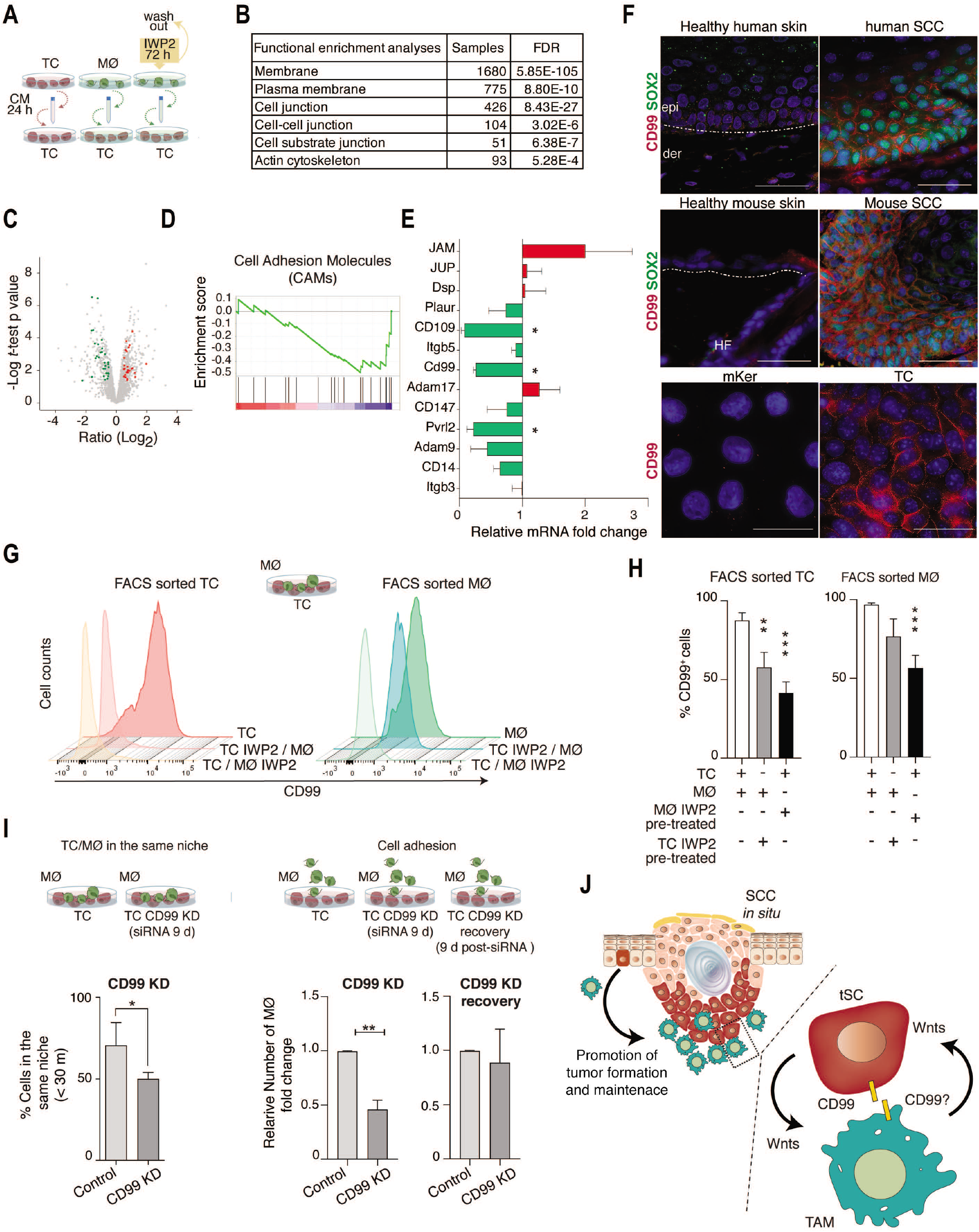
CD99 contributes to the Wnt – mediated cancer cell – macrophage interaction. **(A)** Protocol used to stimulate TC with TC- or MØ-conditioned media (CM) harvested from cells pretreated with IWP-2 or vehicle. **(B)** Functional enrichment analysis showing the top significant cellular component categories among all identified membrane proteins. **(C)** Volcano scatter plot representing fold changes in protein expression (Ratio Log2 FC) of TC stimulated with MØ CM under IWP2 vs vehicle culture conditions. Log2 ratio > 0.5 or < −0.5; p-value < 0.05; False discovery rate (FDR) <0.05. Non-significant (gray), downregulated (green), upregulated (red). Two-sample *t*-test with a permutation-based false discovery rate. **(D)** GSEA plot showing the enrichment of the Cell Adhesion Molecules (CAMs) category. p-value < 0.059, FDR 0.53. **(E)** Relative mRNA expression fold changes of candidate molecules in TC stimulated with MØ CM under IWP2 vs vehicle culture conditions (n = 3). Downregulated (green), upregulated (red). **(F)** Immunostaining for SOX2 (green) and CD99 (red) in human and mouse healthy skin and hSCC and mSCC. Scale bar 50 μm. (Top and Middle). Immunostaining for CD99 (red) in mouse keratinocytes (mker) and TC. Scale bar 20 μm. (Botton). DAPI nuclear staining is represented in blue. **(G)** Histograms of the number of CD99^+^TC and CD99^+^MØ grown in co-culture, comparing cells treated with vehicle versus IWP2 (n = 3). **(H)** Quantification of CD99 membrane expression levels in FACS-sorted TC (Left) or MØ (Right) from TC/MØ co-cultures grown under the indicated IWP2 pre-treatment and vehicle conditions (n = 3). **(I)** Experimental approaches used to analyze the requirement of CD99 expression in TC for the association between TC and MØ (top). Quantification of MØ in the TC niche (<30 μm) in control and CD99 KD TC (Left) and FACS quantification of adhered MØ to TC (Right) in control and CD99 KD TC (n = 3). **(J)** Model of the Wnt-mediated SCC initiation and sustenance of skin cancer stemness by a macrophage niche. epi, epidermis; der, dermis; HF, hair follicle. n represents the number of independent experiments. **(E)** Two-way Anova. (**H)**, One-way Anova. **(I)**. Unpaired two-tailed Student’s *t*-test. All error bars represent s.e.m. *P < 0.05; **P < 0.01; ***P < 0.001.

### CD99 requirement for the tSC-TAM adhesion

From the differentially expressed membrane proteins that could potentially mediate the Wnt-dependent TC-MØ interaction we selected CD99 due to its prior implications in mediating homotypic and heterotypic adhesions, and immune cell migration^47–51^. CD99 is a transmembrane protein that exerts different roles in tumorigenesis in a tissue dependent manner, functioning either as tumor suppressor or oncogene^45,46^. In SCC, however, the role of CD99 is not known. We found CD99 highly expressed in the cell boundaries of SOX2^+^ tSC in both hSCC and mSCC (Fig. 5F), which suggests an oncogenic function for CD99 in skin. In healthy skin, CD99 expression was nearly absent compared to tumors (Fig. 5F), but in agreement with prior findings^52^, it was expressed in basal epidermal cells, suggesting a link with cell proliferation capacity. In culture, CD99 was also highly expressed in TC compared to primary mouse keratinocytes (Fig. 5F), and the expression of CD99 in both TC and MØ required Wnts (Fig. S8A and S8B). In co-culture, the CD99 mRNA and membrane protein expression levels were reduced in both TC and MØ under IWP2 pre-treatment conditions (Fig. 5G, H). These results revealed that Wnts are required for the expression of CD99 in skin cancer cells and macrophages, and a Wnt signaling loop between skin cancer cells and macrophages regulates the expression of CD99 in both populations.

To determine if CD99 mediates the TC and MØ association, we transiently knocked-down CD99 in TC (CD99 KD). RT-PCR and FACS analyses of CD99 KD TC cells showed a 60% and an 83% reduction on CD99 mRNA and membrane protein levels, respectively (Fig. S8C and S8D). The reduction of CD99 expression in TC led to a cell-autonomous decrease in the expression of the Wnt/β-catenin targets Cyclin D1 and Axin2, suggesting a feedback loop between CD99 expression and Wnt signaling (Fig. S8E). We also observed reductions in cell proliferation, TC colony formation, and active MAPK levels, with an increase in epidermal differentiation and apoptosis (Fig. S8F-J). These *in vitro* results suggest an oncogenic role for CD99 in skin cancer cells, sustaining cell proliferation and preventing cell differentiation and cell death. To mimic the tSC-MØ niche, we co-cultured again TC with MØ. Interestingly, the reductions of CD99 expression in CD99 KD TC were sufficient to prevent the formation of heterotypic colonies with MØ compared to controls (Fig. 5I). Also, the heterotypic cell adhesion of MØ to CD99 KD TC decreased (Fig. 5I and Fig. S8K), and the recovery of TC CD99 expression levels rescued the effect. These results support a role for CD99 in sustaining TC proliferation and in mediating TC-MØ heterotypic interactions within the same niche.

## Discussion

In summary, our results show that SCC formation and maintenance are governed by a Wnt-mediated tSCs – TAMs crosstalk, expanding the cellular types that shape the skin tSC microenvironment and control SCC stemness, including the perivascular niche^7,53^ and the tSC-driven evasion of cytotoxic T cells responses^54^. Skin tSC shape their TAMs milieu through Wnts and, in turn, TAM-derived Wnts sustain tSC features for the initiation and the maintenance of tumors (Fig. 5J). Our results also define that CD99 expression in cancer cells contributes to holding the tSC-TAM heterotypic interactions and cancer cell proliferation in a Wnt dependent manner (Fig. 5H). As we learn more about how genetic-deregulated skin tSC pathways lead to TAMs recruitment to the tSC niche and prompt tumor initiated cells to form tumors, it should become clearer how the correction of abnormal growth of mutated cells by normal epidermal cells^11^ is overruled by TAMs Wnts. Given the roles of TAMs in the malignancy of other tumors^55–57^, it is also tempting to speculate a role for TAMs Wnts in the acquisition of tSC invasive characteristics, which has recently been connected to the recruitment of macrophage expressing TGFβ to the tSC niche of SCC *in situ*^18^. This event can be potentially accompanied with a decline in CD99 expression as it has been linked to the transition from local to invasive neoplasms^46^. With our findings, it will be relevant to investigate their potential therapeutic value for immunoprevention against SCC.

## Methods

No statistical methods were used to predetermine sample size.

### In situ RNA analysis

Double in situ RNA hybridization analyses (RNAscope 2.5 HD Duplex Assay, Advanced Cell Diagnostics (ACD), Hayward, CA) were performed on hSCC tissue arrays (SK801 and SK801c TMA, US Biomax)^50^. Wnt ligand expression was screened using the following RNAscope target probes: Wnt2b (Hs-WNT2b-C1, ACD, #453361), Wnt3a (Hs-WNT3a-C1, ACD, #429431), Wnt7a (Hs-WNT7a-C1, ACD, #408231), Wnt7b (Hs-WNT7b-C1, ACD, #421561), Wnt9a (Hs-WNT9a-C1, ACD, #457931) and Wnt10a (Hs-WNT10a-C1, ACD, #429469). CD68 (Hs-CD68-C2, ACD, #560591) was used as a pan-macrophage marker^15^ and Sox2 (Hs-SOX2-C1, RNAscope, ACD, #400871) as a tumor-initiating cell marker^13,14^. The total amount of CD68^+^ cells and double CD68^+^/Wnt^+^ cells within the tumor area were manually calculated. The distance between CD68^+^ cells and Sox2^+^ cells was measured using the ImageJ software. The RNAscope signal per cell (visible at 20–40x magnification) was categorized into 4 grades according to the following scoring guidelines: score 0, no staining or less than 1 dot for every 10 cells; score 1 (+), 1–3 dots per cell; score 2 (++), 4–10 dots per cell with very few dot clusters; score 3 (+++), >10 dots per cell.

### Mouse models

Mice were housed in the AAALAC-accredited pathogen-free (SPF) animal house of the Spanish National Cancer Research Centre (CNIO), Madrid. All animal experimental protocols were approved by the IACUC and the Animal Experimental Ethics Committee of the Carlos III Health Institute (Madrid, Spain), in accordance with National and European regulations. The following commercially available mouse lined were used: CD68CreERT2 (Tg(Cd68-creERT2)31.11.ICS, Institut Clinique de la Souris), K15CrePR1 (Tg(Krt1-15-cre/PGR)22Cot/J, Jackson Labs #005249)^51^, LysMCre (B6.129P2-Lyz2tm1(cre)Ifo/J, Jackson Labs,#004781)^52^, Wls^lox/flox^ (129S-Wlstm1.1Lan/J, Jackson Labs, #012888)^24^, TCF/Lef1-EGFP (Tg(TCF/Lef1-HIST1H2BB/EGFP)61Hadj/J, Jackson Labs, #013752)^53^ and K15GFP (Krt1-15-EGFP2Cot, Jackson Labs, #005244)^51^. Mice were backcrossed to C57BL/6J mice. Both males and females were used in this study with blind inclusion and housed with a light/dark cycle in a temperature-controlled room (22 ± 1°C) with water and food ad libitum.

### Skin tumorigenesis and tumor regression

Mice were treated with DMBA (7,12-dimethylbenz[a]-anthracene) (Sigma-Aldrich) and TPA (12-O-tetradecanoyl phorbol-13-acetate) (Sigma-Aldrich) as previously described^20,21^. Briefly, DMBA (200 μL of 0.25 mg/mL in acetone) was applied once onto the shaved back skin of 6-week-old mice followed by 4 d of TPA treatment (200 μL of 0.02 mg/mL in acetone). TPA was then applied twice weekly for 25 or 30 weeks. 5 weeks-old K15CrePR,Wls^lox/lox^ mice were treated with mifepristone (Sigma-Aldrich) (10 mg/mL dissolved in sunflower seed oil, 2 mg per day) via i.p. injections during 4 consecutive days^27^ immediately before of the DMBA-TPA protocol. Through the duration of the DMBA-TPA treatment, 429 ng/mL mifepristone were administered orally in the drinking water^54^ until the end of the experiment. 5 weeks-old Cd68CreERT2;Wls^lox/lox^ mice were treated with tamoxifen (Sigma-Aldrich) (10 mg/mL dissolved in sunflower seed oil, 2 mg per day) via i.p. during 4 consecutive days before the DMBA-TPA treatment, and through the duration of the experiments, tamoxifen was administered orally in the pellet food. Mice were sacrificed at 25 or 30 weeks after the initiation of the treatments or when tumors reached 1cm or mice presented any signs of distress. To perform the tumor regression experiments, both control and sample groups were treated with DMBA-TPA as previously described, until the appearance of 5 mm tumors, before the administration of tamoxifen or mifepristone for subsequent 8 weeks. Skin tumors were measured weekly using a precision caliber to evaluate lesion size changes (> 0.1mm).

### Histology, immunostaining and image analyses

Skins were embedded in OCT compound (Thermofisher) and stored at −80°C. 10 μm skin sections were fixed in 4% paraformaldehyde (PFA) (Electron Microscopy Sciences) for 10 min at room temperature (RT), then washed in PBS three times and incubated for 1 h in blocking solution (1% BSA, 0.3% Triton X-100, 5% fetal bovine serum and 1% gelatin from cold water fish skin in PBS). Primary antibodies were incubated overnight at 4°C (See Supplementary Table S1). Sections were rinsed three times in PBS and incubated with secondary antibodies during 1h at RT (See Supplementary Table S1). Nuclei were stained with DAPI (Sigma-Aldrich,). Slides were mounted using Mowiol (Sigma-Aldrich) supplemented with 2.5% DABCO (Sigma-Aldrich).

For histological analysis, 6 μm paraffin-embedded sections were deparaffinized, rehydrated and stained with hematoxylin and eosin. For immunostaining, sections were incubated in 10 mM Citrate buffer (pH 6) at 98°C for 15 min for antigen retrieval and incubated in blocking buffer or the MOM Basic kit (Vector Labs) and stained with antibodies as previously described.

Cells cultured on coverslips were fixed in 4% PFA (Electron Microscopy Sciences, Cat. N. 15710) in PBS for 10 min followed by incubation in blocking solution for 1 h. Cells were incubated with antibodies as mentioned above.

Individual images were captured using an Eclipse 90i fluorescent microscope (Nikon Instruments adapted with a DS-QiMC digital camera and the NIS-elements acquisition software) or a Leica TCS SP5 confocal microscope (Leica microsystems). Whole slide scans were acquired with an AxioScan Z1 slide scanner (Zeiss). Either the Zen Blue software (Zeiss) or the Fiji version of the ImageJ software (NIH) were used for quantification of positive cells. 8-10 images per sample were obtained and used for analysis and quantification.

### Cell lines and treatments

The mouse RAW 264.7 cell line^31^ (ATCC, TIB-71) was grown in DMEM supplemented with 10% fetal bovine serum, and 1% antibiotics at 37°C in a humidified 5% CO2 atmosphere. The mouse HACA4 cell line^30,55^ was grown in DMEM/F12 medium supplemented with 10% fetal bovine serum, and antibiotics at 37°C in a humidified 5% CO2 atmosphere. For cells growing in co-cultures, 3×10^5^ HaCa4 and RAW 264.7 cells were each plated on 10 μg/mL fibronectin coated coverslips for 24 h.

Primary mouse keratinocytes were isolated from newborn mouse back-skin, as previously described^56^, and grown in 0.05 mM Ca^2+^ MEM medium supplemented with 15% calcium-chelated FBS.

IWP2 treatment. Cells were grown at 30% confluence before the addition of 10 μM IWP-2 (Sigma Aldrich). Cells were incubated at 37°C in a humidified 5% CO2 atmosphere for 72 h.

Treatment with conditioned media. RAW 264.7 grown at 30% confluence were serum starved in 1% FBS DMEM/F12 medium for 24 h. Media were collected, centrifuged and filtered through 40 μm filter to remove cell debris, and added to cancer cell cultures for 24 h.

### Flow cytometry and cell sorting

Single cell suspensions were obtained from healthy mouse skin or tumors as previously described^57^. Briefly, tissues were minced into small pieces and incubated in 0.25% Trypsin-EDTA solution (Gibco) overnight at 4°C (tumor-initiating cells) or 2 h at RT (myeloid cells and macrophages). Cells were resuspended with 2% calcium chelated FBS, filtered through 70 μm cell strainers and reserved at 4 °C. The remaining tissue on the filter was digested in PBS, 1% BSA, 0.5 mg/ml DNase I (Roche), and 0.5 mg/ml collagenase II and IV (Gibco) for 45 min at 37°C. Single cell-suspensions were obtained via pipette mechanical dissociation, filtered through 40 μm cell strainers, and pooled with the cell suspensions obtained before. Cells were washed in PBS, blocked using the mouse seroblock FcR reagent (CD16/CD32), and stained for FACS analysis in ice-cold PBS, 0.5% BSA, 0.3 mM EDTA using different antibodies (See supplementary Table S1), as follows: Macrophage lineages CD45, MHCII, Ly6C, Ly6G, CD11b, F4/80 antibodies, Myeloid cells CD45, Gr1, CD11b, F4/80, tumor-initiating cells and HFSC CD45, CD34, CD49f and EpCam.

Cultured cells were resuspended and incubated with the Intracellular Fixation & Permeabilization Buffer Set (eBioscience) following the manufacturer’s recommendations and stained for CD34 and SOX2.

FACS analyses were performed using a FACS Canto II (Becton Dickinson), or a FACS ARIA IIu sorter (Becton Dickinson) or InFlux (Cytopeia-BD) for cell sorting. The FlowJo software was used to analyze the data.

### qRT–PCR

RNA extraction from FACS-isolated cells was performed using the Absolutely RNA nanoprep kit (Stratagene) according to the manufacturer’s recommendations. RNA from cells in culture or frozen skin/tumors was isolated using TRIZOL (Sigma-Aldrich) using standard procedures. 1 μg was used for cDNA synthesis using the Maxima First Strand cDNA synthesis kit (Thermo Scientific). Quantitative PCR assays were performed using 1 ng of cDNA as template and the reactions were conducted using the GoTaq qPCR Master Mix (Promega). Real-time RT-PCR was performed on a MasterCycler Ep-Realplex thermal cycler (Eppendorf) or an Applied Biosystems QuantStudio™ 7 Flex Real-Time PCR (Applied Biosystems). β-actin and GAPDH housekeeping genes were used for normalization. The gene-specific primers sets were used at final concentration of 0.2 μM and their sequences are listed in supplementary Table S1. All qRT–PCR assays were performed in triplicate in three independent experiments.

### Transmission electron microscopy

Cells were grown at confluence on Thermanox^TM^ coverslips and fixed with 2% v/v glutaraldehyde in 0.05 M sodium phosphate buffer (pH 7.2). Samples were rinsed three times in 0.15 M Phosphate buffer (pH 7.2) and subsequently post-fixed in 1% w/v OsO4 in 0.12 M sodium Phosphate buffer (pH 7.2) for 2 h. The specimens were dehydrated in graded series of ethanol, transferred to propylene oxide and embedded in Epon according to standard procedures. Following polymerization, the Thermanox^TM^ coverslip was removed. 60 nm thick sections were cut with a Leica UC7 microtome (Leica Microsystems, Wienna, Austria) and collected on copper grids with Formvar supporting membranes, stained with uranyl acetate and lead citrate, and subsequently examined with a Philips CM 100 Transmission EM (Philips, Eindhoven, The Netherlands), operated at an accelerating voltage of 80 kV. Digital images were recorded with an OSIS Veleta digital slow scan 2k x 2k CCD camera and the ITEM software package.

### Proliferation assay

1×10^4^ cells were seeded per well in 24-well plates for 24, 48, 72 and 96 h in medium without antibiotics. Cells were transfected with Lipofectamine-RNAiMAX (13778100, Thermo Fisher Scientific) and re-transfected every 72 h. At the indicated time points, the cells were trypsinized, stained with 1:1 Trypan Blue to discriminate dead cells and counted in a Neubauer chamber.

### Colony formation assay

HaCa4 cells controls and treated were plated at a density of 500 cells/well on 6-well plates for 10 d. Cells were fixed in ice-cold methanol for 10 min and stained with 1% Crystal violet (Sigma-Aldrich) for 15 min.

### Live imaging microscopy

Cells (pre-treated with IWP2 or vehicle) were seeded onto cell culture chambers with and insert (Ibidi). 24 h later the cells were treated with mitomycin C (Roche) for 1 h at 37°C, washed and cultured with either fresh medium, or conditioned media. Bright field images were captured every 10 min for 72 h using an inverted motorized microscope (Leica DMI 6000) coupled with an incubation system to control the temperature and CO2 levels during the course of the experiments. The area covered by each population was measured and quantified using ImageJ software.

### Adhesion assay

0.3 x 10^6^ HaCa4 cells were seeded in 6-well plates at 30% and grown at ~90% confluence. RAW 264.7 macrophage-like cells were pellet by centrifugation and resuspended in DMEM medium. 1.2 x 10^6^ cells/mL were added on top of HaCa4 cell monolayers in a 3:2 ratio. After 2 h, the non-adherent RAW 264.7 cells were gently rinsed away by washing twice plates with PBS. The remaining adhered cells were harvested, incubated with the mouse seroblock FcR reagent (CD16/CD32), and stained using CD45, CD34, CD11b, F4/80 antibodies (See supplementary Table S2), and analyzed by FACS as previously described.

### siRNA transfection

The CD99 knockdown in HaCa4 cells was achieved using the ON-TARGET plus SMART pool siRNA targeting mouse CD99 (J-165109-01-0005 and J-165109-09-0005 ON-TARGET plus, Dharmacon), and siRNA scramble controls (AM4621, Thermofisher). siRNAs were transfected using Lipofectamine-RNAiMAX (13778100, Thermo Fisher Scientific) according to the manufacturer’s instructions.

### Immunoblot

Cells were lysed in RIPA buffer (40 mM Tris-HCl, pH 7.4, 150 mM NaCl, 1% NP-40, 0.5% DOC, 0.2% SDS, 2 mM EDTA) complemented with protease inhibitors. The protein concentration was quantified using the BCA protein assay kit (Pierce). SDS-PAGE was performed using standard procedures.

### Proteomic analyses

Membrane enrichment was performed as previously reported^58^. Briefly, cells were lysed in 500 μl of lysis buffer (8 mL of cold PBS and 2 mL of 10% Triton X-114 supplemented with Halt Protease inhibitor (1:100) and Bezonase (Merck) (1:1000)). The pellets were vortexed and left overnight at 4°C on a rotary shaker. Samples were centrifuged at 20 000 x g for 30 min at 4°C. Supernatants were collected and placed in a thermal mixer at 37°C at 700 rpm for 30 min and centrifuged at 5000 x g for 30 min at 25°C, to separate detergent and aqueous phases. The aqueous top layer was removed, and the bottom detergent phase was resuspended in 0.8 mL of cold PBS. The samples were placed in a thermal block at 37°C and 900 rpm for 15 min followed by 15 min centrifugation at 5000 x g and 25°C. This process was repeated three times in order to wash the detergent phase and remove contaminants. Proteins were extracted by acetone precipitation: 600 μl of acetone were added to the detergent phase and incubated overnight at −20° C. Samples were spun at 5 000 x g for 30 min at 4°C. Supernatants were removed and the pellets were resuspended in 200 μl of 8 M urea in 100 mM Tris pH8. Protein concentrations were measured with the Qubit® Protein Assay Kit.

20 μg of each sample were digested by means of the standard FASP protocol. Briefly, proteins were reduced (15 mM TCEP, 30 min, RT), alkylated (50 mM CAA, 20 min in the dark, RT) and sequentially digested with Lys-C (Wako) (protein:enzyme ratio 1:100, o/n at RT) and trypsin (Promega) (protein:enzyme ratio 1:100, 6 h at 37 °C). Resulting peptides were desalted using SCX stage-tips and dissolved in 30 μL of 0.5% formic acid.

### Mass spectrometry

LC-MS/MS was conducted by coupling a nanoLC-Ultra 1D+ system (Eksigent) to an LTQ Orbitrap Velos mass spectrometer (Thermo Fisher Scientific) via a Nanospray Flex source (Thermo Fisher Scientific). Peptides were loaded into a trap column (NS-MP-10 BioSphere C18 5 μm, 20 mm length, Nanoseparations) for 10 min at a flow rate of 2.5 μl/min in 0.1% FA. Peptides were transferred to an analytical column (ReproSil Pur C18-AQ 2.4 μm, 500 mm length and 0.075 mm ID) and separated in buffer A (4% ACN, 0.1% FA; buffer B: 100% ACN, 0.1% FA) at a flow rate of 250 nL/min. The gradient used was: 0-2 min 2-6% B, 2-103 min 6-30%B, 103.5-113.5 min 98% B, 114-120 min 2% B. The peptides were electro sprayed (1.4 kV) into the mass spectrometer with a PicoTip emitter (360/20 Tube OD/ID μm, tip ID 10 μm), at a heated capillary temperature of 325 °C and S-Lens RF level of 60%. The mass spectrometer was operated in a data-dependent mode, with an automatic switch between MS and MS/MS scans using a top 15 method (threshold signal ≥ 800 counts and dynamic exclusion of 60 sec). MS spectra (350-1500 m/z) were acquired in the Orbitrap with a resolution of 60,000 FWHM (400 m/z). Peptides were isolated using a 1.5 Th window and fragmented using collision induced dissociation (CID) with linear ion trap read out at a NCE of 35% (0.25 Q-value and 10 ms activation time). The ion target values were 1E6 for MS (500 ms max injection time) and 5000 for MS/MS (100 ms max injection time).

### Protein data analysis

Raw files were processed with MaxQuant (v 1.6.2.3) using the standard settings against a mouse protein database (SwissProt/TrEMBL, 53449 sequences) supplemented with contaminants. Label-free quantification was done with matches between runs (match window of 0.7 min and alignment window of 20 min). Carbamidomethylation of cysteines was set as a fixed modification whereas oxidation of methionine and protein N-term acetylation were set as variable modifications. The minimal peptide length was set to 7 amino acids and a maximum of two tryptic missed cleavages were allowed. Results were filtered at 0.01 FDR (peptide and protein level).

Afterwards, the “proteinGroups.txt” file was loaded in Prostar (v1.18.2)^59^ using the LFQ intensity values for further statistical analysis. Briefly, proteins with less than three valid values in at least one experimental condition were filtered out. Then, missing values were imputed using the algorithms SLSA^60^for partially observed values and DetQuantile for values missing on an entire condition. Differential analysis was done using the empirical Bayes statistics Limma. Proteins with a p value < 0.05 and a log2 ratio >0.5 or <-0.5 were defined as regulated. The FDR was estimated to be below 5%.

### Statistical analysis

All quantitative data are presented as mean ± SEM (standard error of the mean). Results are representative of at least three independent experiments conducted in triplicates. Comparisons of normally distributed datasets were analyzed using unpaired two-tailed Student’s *t*-test. Non-normally distributed datasets were analyzed using non-parametric tests. Comparisons involving three or more groups were conducted using the ANOVA test. For all statistical analysis, a confidence level of p ≤ 0.05 was considered statistically significant. Experiments involving images were conducted in triplicates of at least three different experiments, and representative images are shown in the figures. For animal experiments, animals were excluded from the analyses in they died or had to be sacrificed following humane end-point criteria. Statistical analyses were done using GraphPad 8 Software.

### Data availability

Data of the quantitative proteomic profiling of membrane-associated proteins has been deposited in the PRIDE-ProteomeXchange database, accession number PXD019989. All other data are included in the Source data and Supplementary information and provided with the paper. Captured microscope images, correspondence and requests for materials should be addressed to the corresponding author (M.PM).

## Acknowledgments

This work was conducted in two institutions. We thank both our CNIO and University of Copenhagen colleagues for their support and suggestions over the course of the project, in particular at Erwin Wagner’s lab and Martin W. Berchtold’s, Lotte Pedersen’s and Søren Christensen’s labs, and also Arturo Mora-Rioja for his comments on the manuscript. We also thank the following facilities: the CNIO mouse facility, the Animal Units of the University of Copenhagen, the CNIO and Biotech Research & Innovation Centre (BRIC) Flow Cytometry facilities, the CNIO Confocal Microscopy Unit, the Core Facility for Integrated Microscopy of the University of Copenhagen, and the CNIO Proteomics Unit for technical support.

## Authors Contributions

S.F. experimental design and analyses, J.C. and A.MS. experimental analyses, E.Z and J.M. proteomic analyses, D.M. confocal microscopy, D.C. samples and experimental analyses, R.L. histopathological analyses, M.PM. study conception, supervision and funding. S.F. and M.P-M. manuscript writing with input from all authors.

## Additional Information

Supplementary Figs 1–8, Supplementary Movie 1, Supplementary Tables 1–2 are provided with the paper. Correspondence and requests for materials should be addressed to the corresponding author (M.PM).

**Figure S1.**
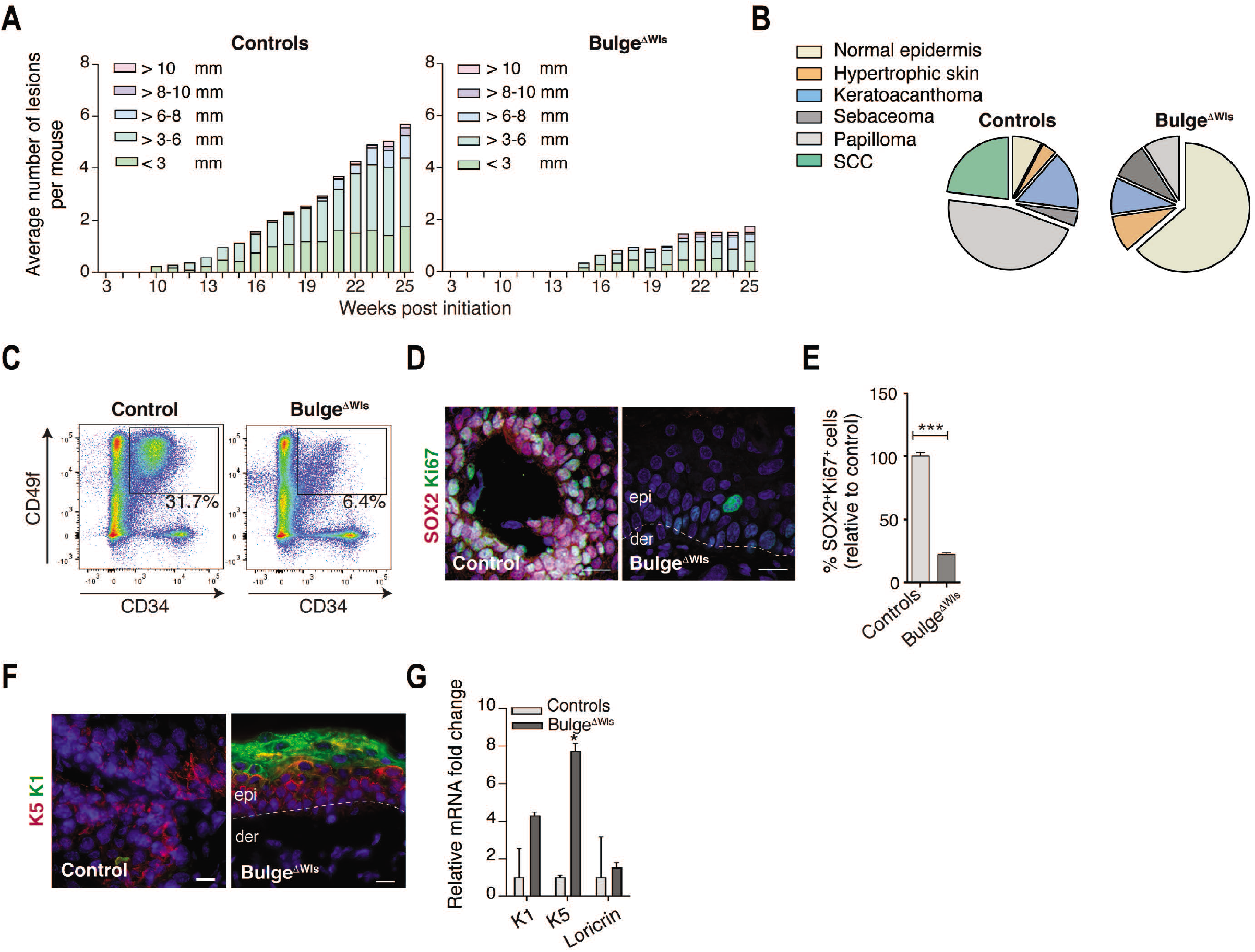
HFSC-derived Wnts are required for skin tumor initiation by promoting tSCs proliferation and preventing tSC differentiation. **(A)** Number of tumors and tumor size over the time course of tumor formation in Bulge^ΔWls^ (n = 17) control (n = 34) mice. **(B)** Pie charts depicting the histological classification of skin lesions in Bulge^ΔWls^ (n = 11) mice and control (n = 26) mice. Tumors were analyzed at the last time point of tumor development. **(C)** Representative FACS plots of isolated CD34^+^CD49f^+^ tSC from skin lesions of Bulge^ΔWls^ (n = 6) mice and control (n=7) mice, respectively. **(D)** Immunostaining for SOX2 (red) and Ki67 (green) in skin sections of control (Left) and Bulge^ΔWls^ mice (Right). **(E)** Quantification of immunostained SOX2^+^Ki67^+^ cells in skin sections of Bulge^ΔWls^ (n = 7) mice and control (n= 12) mice, respectively. **(F)** Immunostaining for K5 (red) and K1 (green) in skin sections of control (Left) and Bulge^ΔWls^ mice (Right). **(G)** qRT–PCR analysis of K5, K1 and Loricrin in total skin of Bulge^ΔWls^ (n = 4 mice) mice and control (n= 4) mice, respectively. epi, epidermis; der, dermis. DAPI nuclear staining is represented in blue. All scale bars represent 20 μm. **(E)** and **(G)** Mann-Whitney test. All error bars represent s.e.m. * P<0.05; ***P < 0.001.

**Figure S2.**
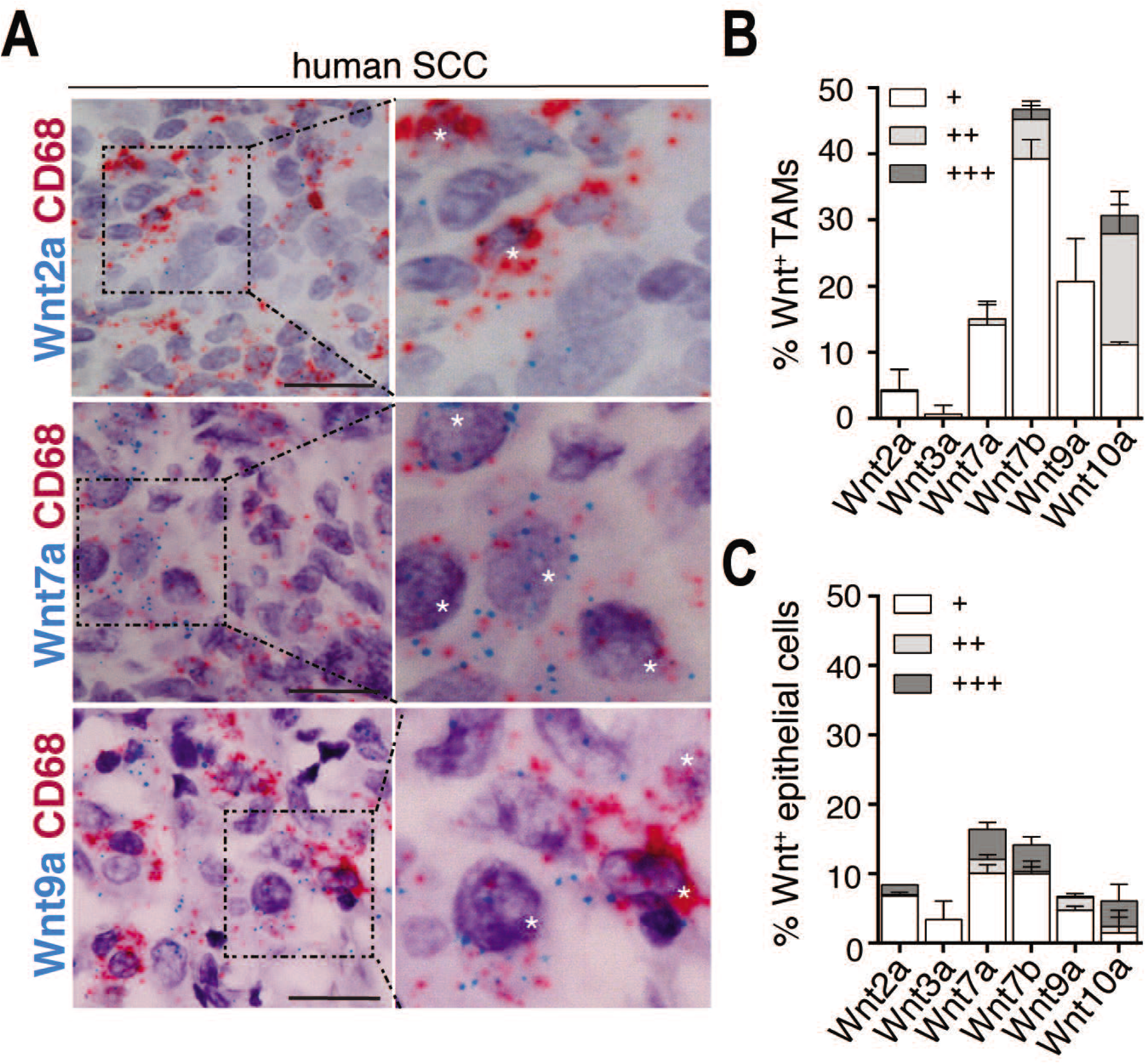
Expression of Wnt ligands by macrophages and cancer epithelial cells in hSCC. **(A)** RNA *in situ* analysis for CD68 (TAMs, red) and Wnt2a (blue, top), Wnt7a (blue, middle) and Wnt9a (blue, bottom) in hSCC. **(B)** Percentage of TAMs in hSCC expressing the indicated Wnts mRNA (score +:1-3 dots; ++:4-10 dots; +++: >10 dots) (n = 42 tumors). **(C)** Quantification of the mRNA expression of the indicated Wnts in epithelial cells (score +: 1–3 dots; ++:4–10 dots; +++: >10 dots) in hSCC (n = 29). Scale bar 20 μm. Insets are magnified views. Asterisks denote positive stained TAMs. Data represent mean ± s.e.m.

**Figure S3.**
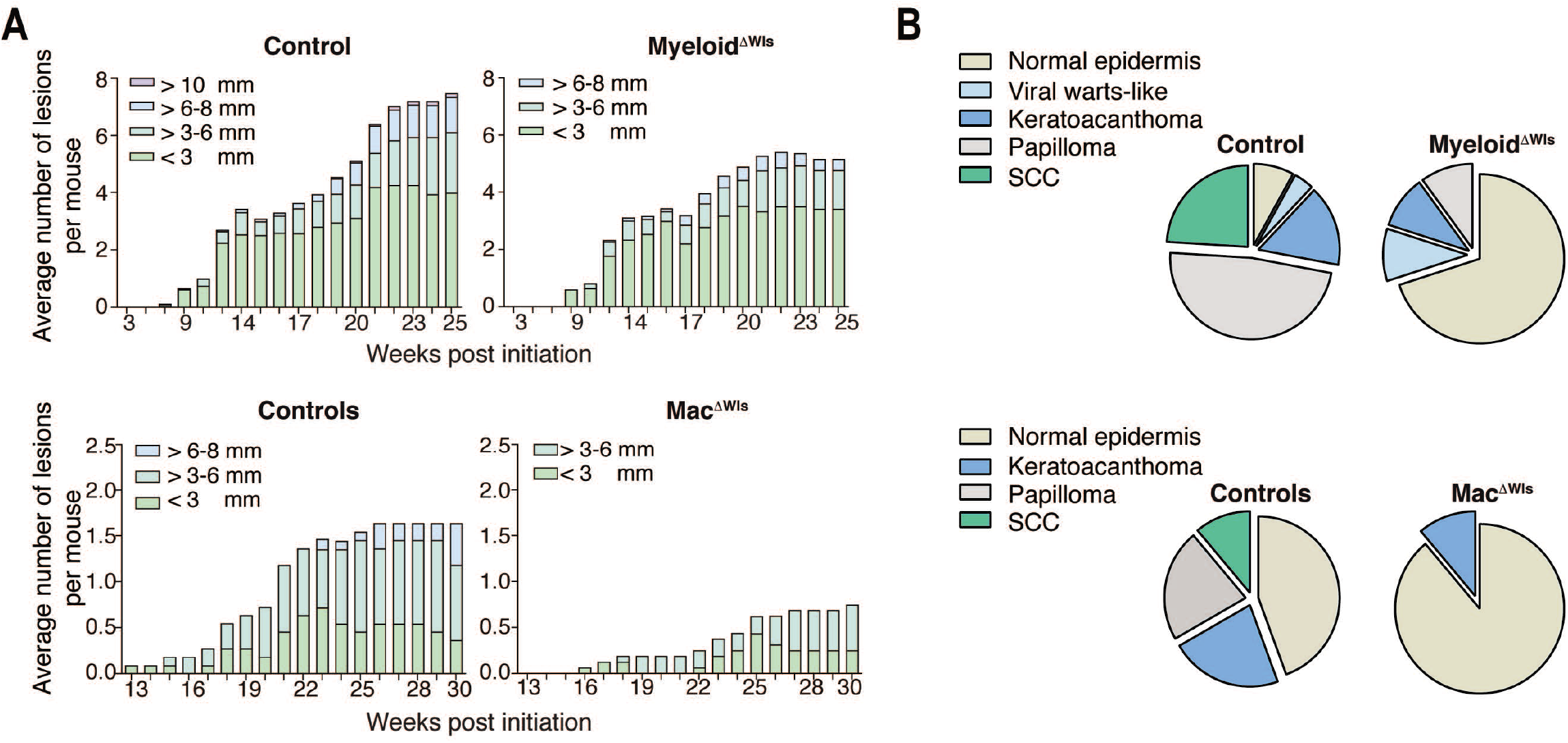
Myeloid- and macrophage- derived Wnts promote skin tumor formation. **(A)** Number of tumors and tumor size over the time course of tumor formation in Myeloid^ΔWls^ (n = 24) (Top) and Mac^ΔWls^ (n = 16) (bottom) mice and control (n = 24, 11) mice, respectively. **(B)** Pie charts depicting the histological classification of skin lesions in Myeloid^ΔWls^ (n = 10) mice (Top) and Mac^ΔWls^ (n = 9) (Bottom) mice and control (n = 25, 9) mice, respectively, at the last time point of tumor development.

**Figure S4.**
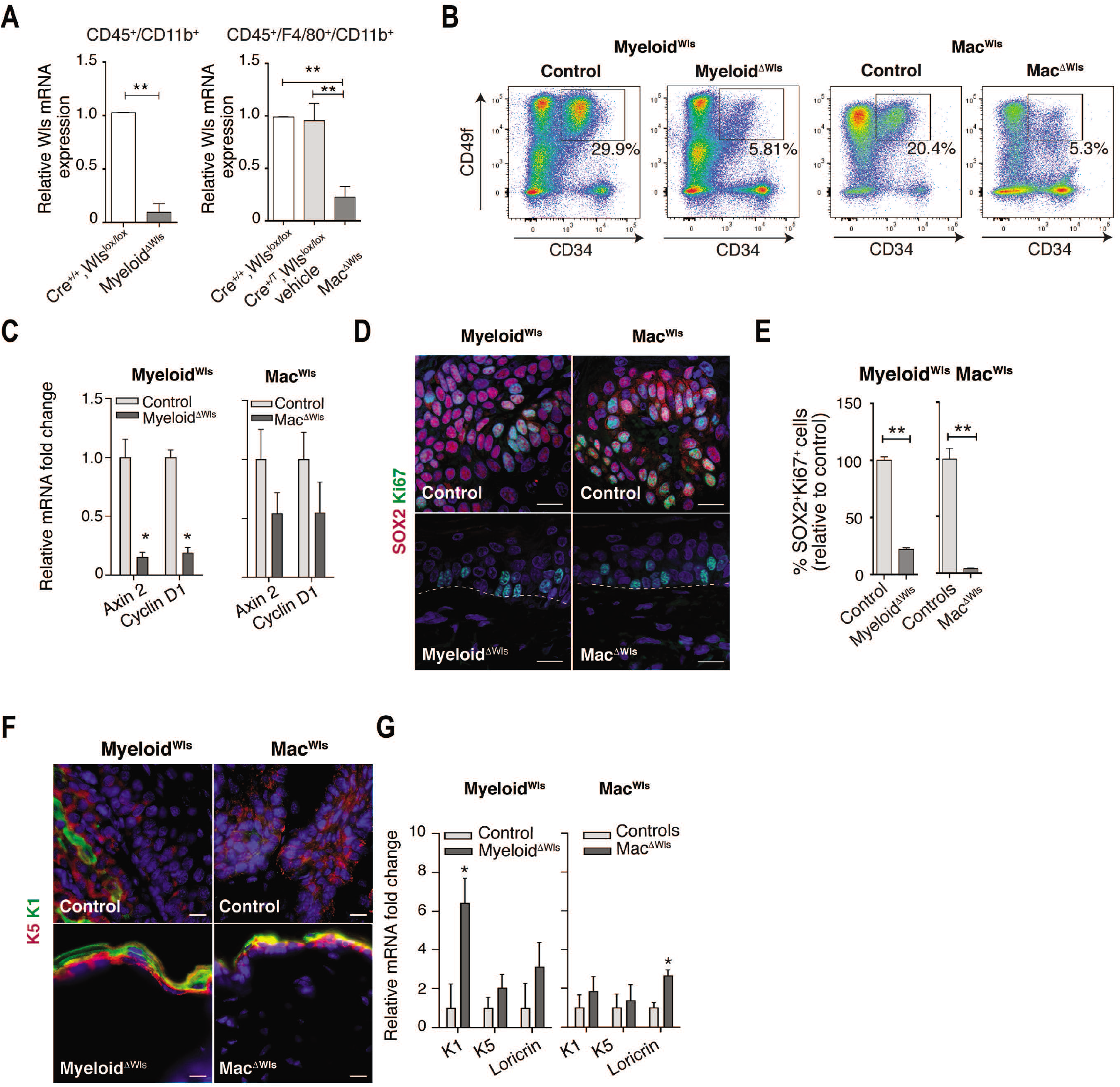
Wnt-expressing myeloid cells and macrophages foster tSC proliferation and prevent tSC differentiation. **(A)** qRT–PCR analysis of Wls mRNA expression in FACS-isolated CD45^+^CD11b^+^ (Myeloid^Wls^, n= 3), and CD45^+^CD11b^+^F4/80^+^ (Mac^Wls^, n= 5) cells from the back skin of the indicated mouse lines and controls (n = 4, 12) mice, respectively. **(B)** Representative FACS plots of isolated CD34^+^CD49f^+^ tSC from skin lesions of Myeloid^ΔWls^ (n = 12) and Mac^ΔWls^ (n =6) mice and control (n=15,8) mice, respectively. **(C)** qRT–PCR analysis of the Wnt/β-catenin targets Axin2 and CyclinD1 expression in total skin of Myeloid^ΔWls^ (n = 4) and Mac^ΔWls^ (n = 4) mice and control (n= 4,4) mice, respectively. **(D)** Immunostaining for SOX2 (red) and Ki67 (green) in skin sections of control and Myeloid^ΔWls^ (Top) and Mac^ΔWls^ mice (Bottom). **(E)** Quantification of immunostained SOX2^+^Ki67^+^ cells in skin sections of Myeloid^ΔWls^ (n = 8) and Mac^ΔWls^ (n = 6) mice and control (n= 5, 6) mice, respectively. **(F)** Immunostaining for K5 (red) and K1 (green) in skin sections of control (Top) and Myeloid^ΔWls^ and Mac^ΔWls^ mice (Bottom). **(G)** qRT–PCR analysis of K5, K1 and Loricrin in total skin of Myeloid^ΔWls^ (n = 4 mice) and Mac^ΔWls^ (n = 4 mice) mice and control (n= 4,4) mice, respectively. DAPI nuclear staining is represented in blue. All scale bars represent 20 μm. **(A)** Unpaired two-tailed Student’s *t*-test and One-way Anova **(C)** Two-way Anova. **(E)** and **(G)** Mann-Whitney test. **(G)** All error bars represent s.e.m. * P<0.05; ** P<0.01.

**Figure S5.**
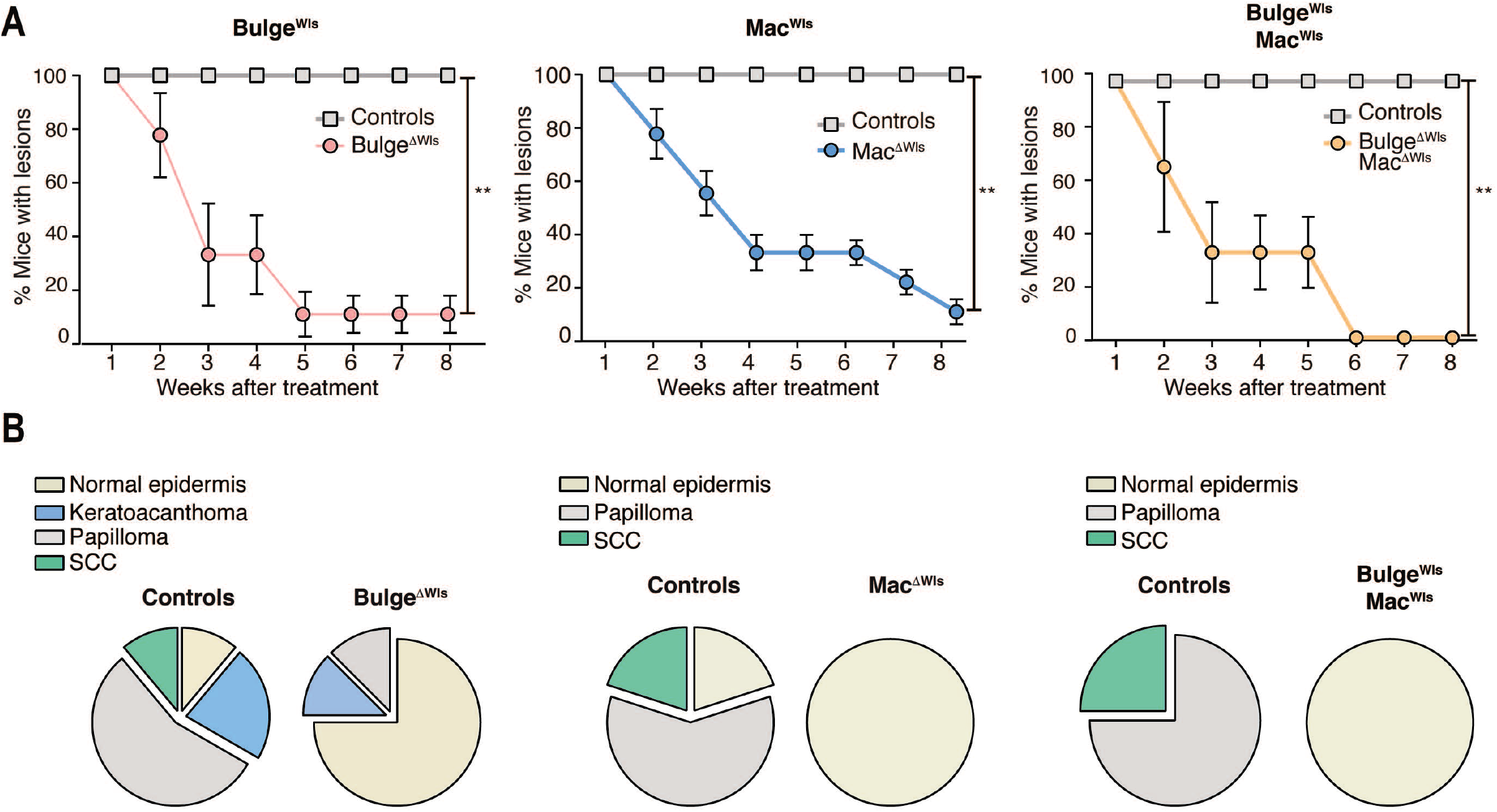
Loss of mature Wnts in either tSCs or TAMs lead to the tumor relapse. **(A)** Percentage of mice presenting skin tumors from the initial time point to 8 weeks of the respective administration of RU467, TAM or RU486+TAM in Bulge^ΔWls^ (n = 9), Mac^ΔWls^ (n = 9), and Bulge^ΔWls^Mac^ΔWls^ (n = 3) mice and control (n = 12, 12, 4) mice, respectively. Unpaired two-tailed Student’s *t*-test. **(B)** Pie charts depicting the histological classification of skin lesions in Bulge^ΔWls^ (n = 8) (left), Mac^ΔWls^ (n = 4) (middle) and Bulge^ΔWls^ Mac^ΔWls^ (n = 3) (right) mice and control (n = 10, 4, 3) mice, respectively, at the last time point of tumor regression. All error bars represented s.e.m. **P < 0.01.

**Figure S6.**
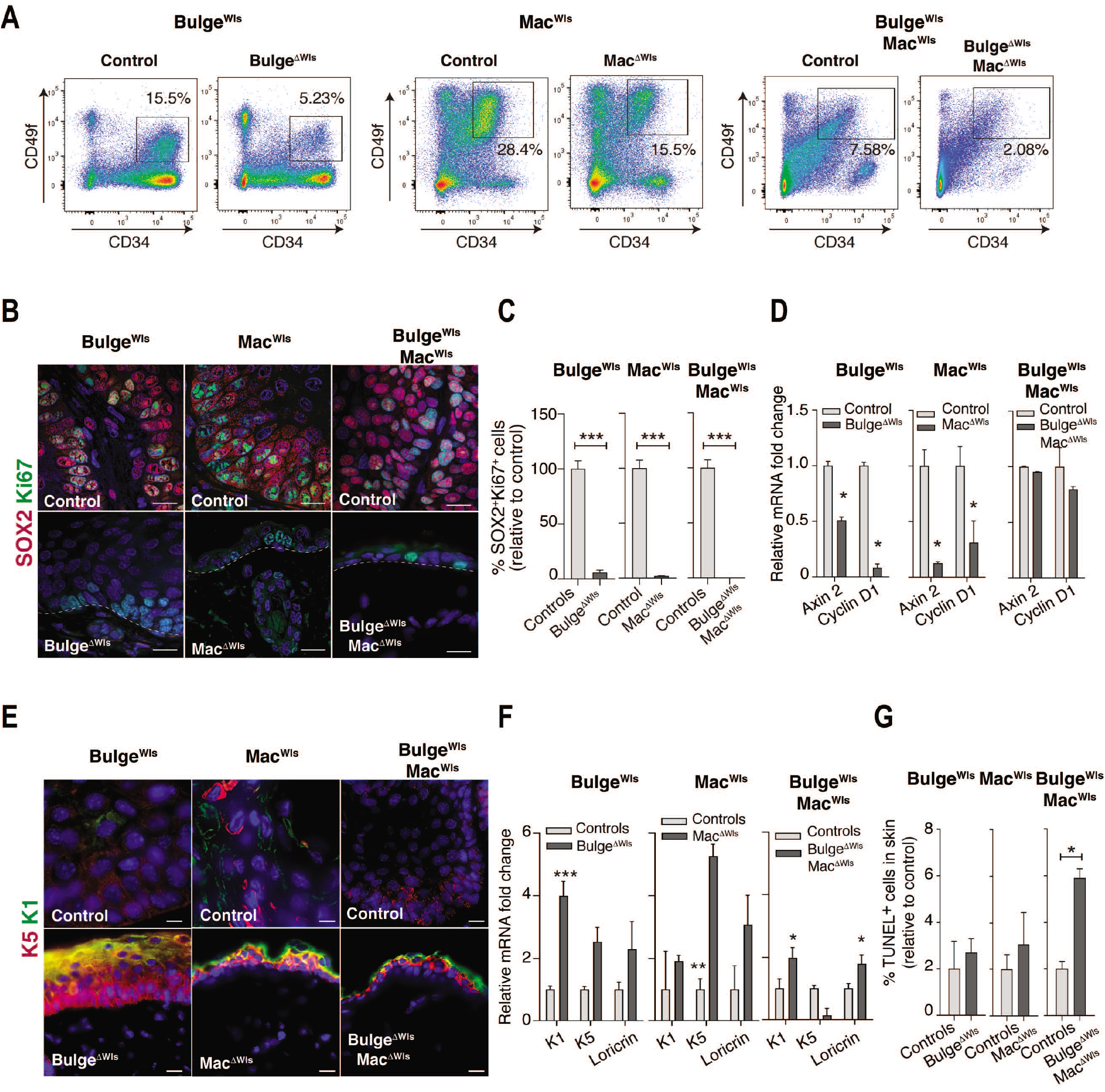
tSC and TAM-derived Wnts prevent tumor relapse by promoting tSC proliferation and precluding epidermal differentiation. **(A)** Representative FACS plots of isolated CD34^+^CD49f^+^ tSC from skin lesions of Bulge^ΔWls^ (n = 6), Mac^ΔWls^ (n = 6) and Bulge^ΔWls^ Mac^ΔWls^ (n = 4) mice and control (n= 8,7,3) mice, respectively. **(B)** Immunostaining for SOX2 (red) and Ki67 (green) in skin sections of control (Top) and Bulge^ΔWls^, Mac^ΔWls^ and Bulge^ΔWls^Mac^ΔWls^ mice (Bottom). **(C)** Quantification of immunostained SOX2^+^Ki67^+^ cells in skin sections of Bulge^ΔWls^ (n = 3), Mac^ΔWls^ (n = 3) and Bulge^ΔWls^Mac^ΔWls^ (n = 3) mice and control (n= 3,3,3) mice, respectively. **(D)** qRT–PCR analysis of the Wnt/β-catenin targets Axin2 and CyclinD1expression in total skin of Bulge^ΔWls^ (n = 4), Mac^ΔWls^ (n = 4) and Bulge^ΔWls^Mac^ΔWls^ (n = 4) mice and control (n= 4,4,4) mice, respectively. **(E)** Immunostaining for K5 (red) and K1 (green) in skin sections of control (top) and Bulge^ΔWls^, Mac^ΔWls^ and Bulge^ΔWls^Mac^ΔWls^ mice (bottom). **(F)** qRT-PCR analysis of K5, K1 and Loricrin in total skin of Bulge^ΔWls^ (n = 4 mice), Mac^ΔWls^ (n = 4 mice) and Bulge^ΔWls^Mac^ΔWls^ (n =4 mice) mice and controls (n= 4,4,4) mice, respectively. **(G)** Quantification of TUNEL^+^ cells in skin sections of Bulge^ΔWls^ (n = 5 mice), Mac^ΔWls^ (n = 3 mice) and Bulge^ΔWls^Mac^ΔWls^ (n = 3 mice) mice and control (n= 3,3,3) mice respectively. DAPI nuclear staining is represented in blue. All scale bars represent 20 μm. Analyses were performed at the at the last time point of tumor regression. **(B)**, **(D)** and **(F)** Mann-Whitney test. **(G)** Unpaired two-tailed Student’s *t-* test with Welsh’s correction. All error bars represent s.e.m. * P<0.05; ** P<0.01, ***P < 0.001.

**Figure S7.**
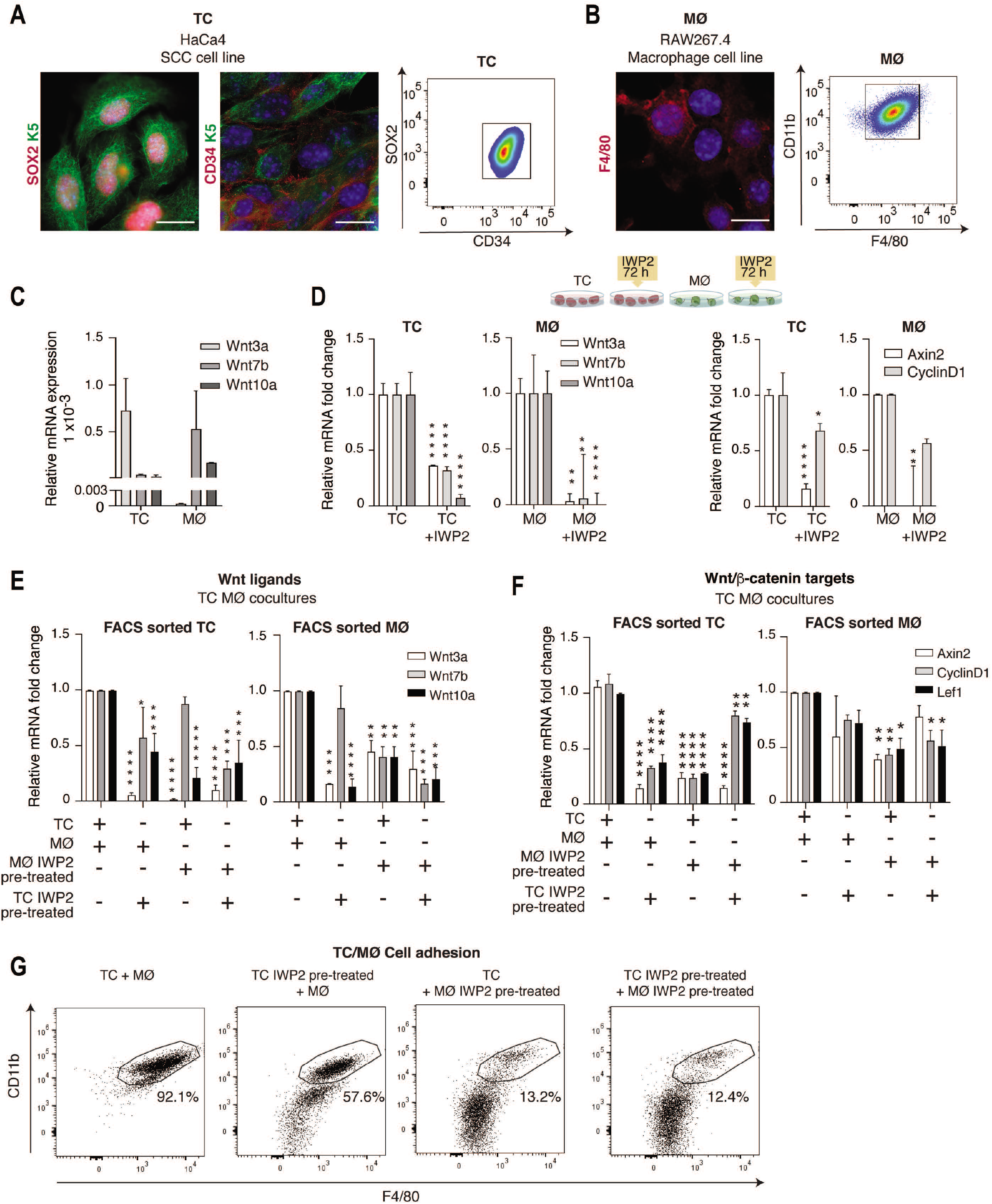
A Wnt feedback loop between cancer cells and macrophages governs Wnt/β-catenin signaling activity through their heterotypic adhesion. **(A)** Immunostaining for K5 (green) and SOX2 (red) and K5 (green) and CD34 (red) in the HaCa4 mouse SCC line (Tumor cells, TC) (Left). Scale bar 10 μm. FACS plot of the expression of SOX2 and CD34 in TC (n = 2) (Right). **(B)** Immunostaining for F4/80 (red) in the RAW 267.4 mouse cell line (macrophages, MØ) (Left). Scale bar 10 μm. FACS plot of the expression of CD11b and F4/80 in MØ (n = 2) (Right). **(C)** qRT–PCR analysis of the expression of Wnt3a, Wnt7b and Wnt10a in TC and MØ (n = 3). **(D)** Protocol used to inhibit the secretion of Wnt ligands in TC and MØ using the IWP2 inhibitor or vehicle control (Top). qRT–PCR analysis of the expression of Wnt3a, Wnt7b and Wnt10a (Left) and the expression of Axin2 and CyclinD1 in TC and MØ treated with IWP2 or vehicle (n = 3) (Right). **(E)** qRT–PCR analysis of the expression of Wnt3a, Wnt7b and Wnt10a in FACS-isolated TC (n = 3) (Left) or MØ (n = 3) (Right) from co-cultures pre-treated with IWP2 or vehicle. **(F)** qRT–PCR analysis of the expression of Axin2, CyclinD1 and Lef1 in FACS-isolated TC (n = 3) (Left) or MØ (n = 3) (Right) from co-cultures pre-treated with IWP2 or vehicle. **(G)** FACS quantification of adhered CD11b^+^F480^+^ MØ to TC under vehicle or IWP2 pre-treatment conditions (n = 3). n represents the number of independent experiments. DAPI nuclear staining is represented in blue. **(C-F)** Two-way Anova. n represents the number of independent experiments. All error bars represent s.e.m. * P<0.05; ** P<0.01, *** P<0.001, **** P<0.0001.

**Figure S8.**
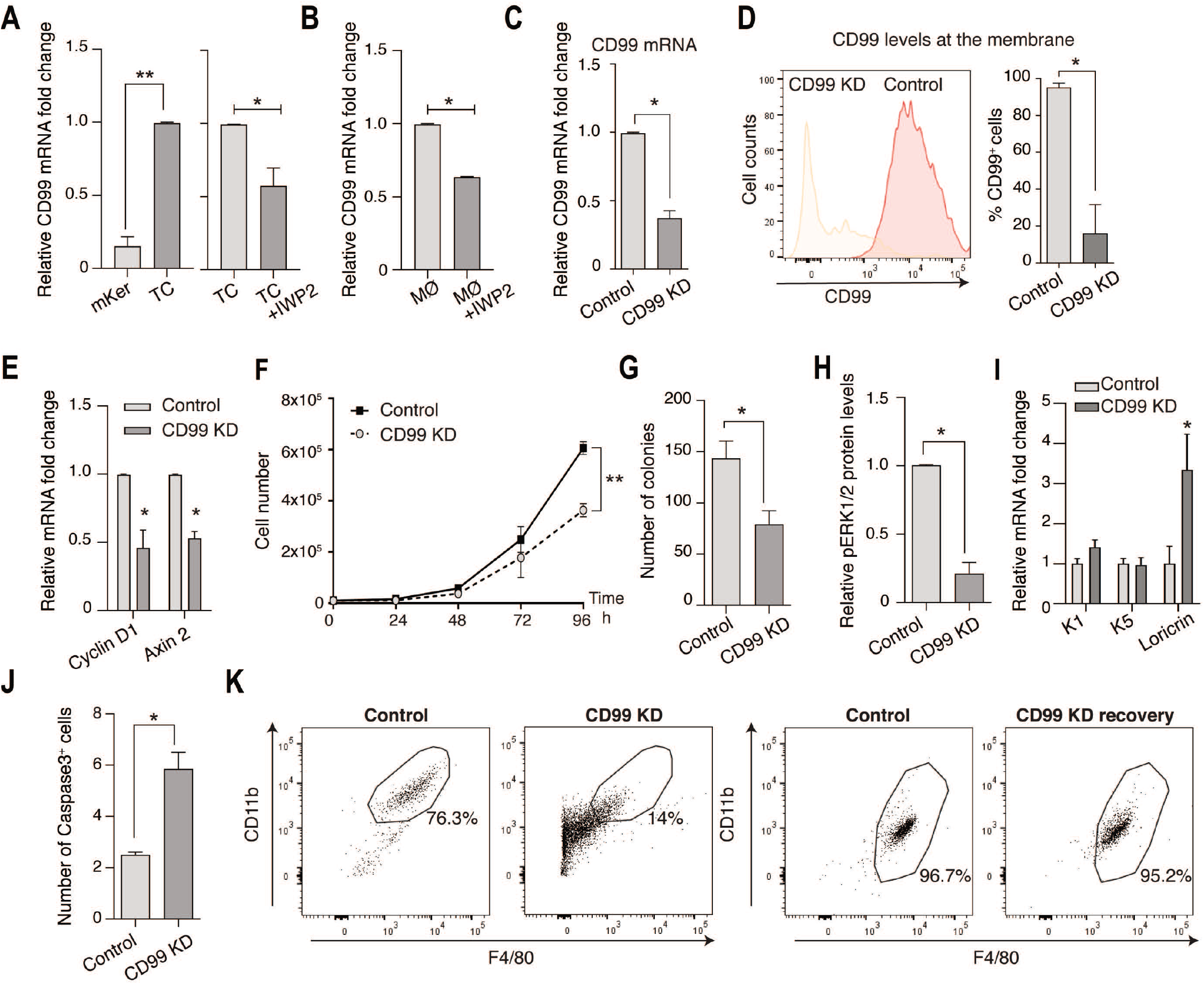
CD99 expression in cancer cells sustains tSC features and cell adhesion to macrophages. **(A)** qRT–PCR analysis of CD99 expression in mKer, TC and TC treated with vehicle or IWP2 (n = 3). **(B)** qRT–PCR analysis of CD99 expression in MØ treated with vehicle or IWP2 (n = 3). **(C)** qRT–PCR analysis of CD99 expression in scramble control and CD99 siRNA knock-down (KD) TC (n = 3). **(D)** FACS analysis of cells expressing CD99 at the membrane in scramble control and CD99 KD TC. Histogram and quantification of CD99^+^cells in scramble control and CD99 KD TC. (n = 3). **(E)** qRT–PCR analysis of the Wnt/β-catenin targets Axin2 and CyclinD1 in scramble control and CD99 KD TC (n = 3). **(F)** Proliferation curves of scramble control and CD99 KD TC (n = 2). **(G)** Quantification of the number of colonies formed by scramble control and CD99 KD TC (n = 3) (left). **(H)** Quantification of phospho-ERK1/2 normalized to total ERK levels in scramble control and CD99 KD TC, assayed by immunoblot (n = 3). **(I)** qRT–PCR analysis of K5, K1 and Loricrin in scramble control and CD99 KD TC (n = 2). **(J)** Quantification of active Caspase 3^+^ cells in scramble control and CD99 KD TC (n = 2). **(K)** FACS quantification of adhered CD11b^+^F480^+^ MØ to scramble control or CD99 KD TC (n = 2). n represents the number of independent experiments. **(A-D)** and **(J)** Unpaired two-tailed Student’s *t*-test. **(E), (F)** and **(I)** Two-way Anova. **(G)** Unpaired two-tailed Student’s *t*-test with Welsh’s correction. All error bars represent s.e.m. *P<0.05, **** P<0.0001.

**Supplementary Table 1.** Commercial antibodies used in the study. The commercial antibodies used in this study are listed together with their source.

| Antibodies | Source | Identifier |
| --- | --- | --- |
| Primary |  |  |
| Goat CD99 | Thermo Fisher | PA5-47551 |
| Rat anti-BrDU | Abcam | Ab6236 |
| Rabbit anti p44/42 MAPK | Cell Signalling | #9102 |
| Rabbit anti 44/42 MAPK | Promega | V8031 |
| Mouse anti-CD68 | Abcam | Ab955 |
| Rabbit anti-cleaved caspase-3 | Cell Signalling | #9661 |
| Rat anti-CD34 | BD Pharmingen | #553731 |
| Rat anti-F4/80 | BioRad | MCA497G |
| Chicken Polyclonal anti-Keratin-5 | Biolegend | #905903 |
| Rabbit anti-Ki67 | Master Diagnostica | MAD-000310QD |
| Rat anti-Sox2 | eBioscience | #14-9811 |
| Rat anti-CD16/CD32 | Becton Dickinson | #553141 |
| Mouse FITC-conjugated anti-BrDu | BioLegend | #364104 |
| Rat FITC-conjugated anti - CD45 | Miltenyi Biotec | #130-102-778 |
| Rat PE-Cy7-conjugated anti-CD45 | eBioscience | #25-0451-81 |
| Rat APC-Cy7-conjugated anti-CD45 | BioLegend | #103116 |
| Rat PERCP-Cy5.5-conjugated anti- CD11b | eBioscience | #45-0112 |
| Rat FITC-conjugated anti-CD11b | eBioscience | #553310 |
| Rat PE-conjugated anti-CD34 | BD Pharmingen | #551387 |
| Rat APC-conjugated anti-CD49f | eBioscience | #17-0495-82 |
| Rat FITC-conjugated anti-Epcam | Biolegend | #118208 |
| Rat APC-eFluor-conjugated anti-F4/80 | eBioscience | #47-4801 |
| Rat PE-Cy7-conjugated anti-F4/80 | Biolegend | #1054780 |
| Rat Alexa 647-conjugated anti-F4/80 | eBioscience | #123122 |
| Rat PE-Cy7-conjugated anti- Gr1 | eBioscience | #25-5931 |
| Rat FITC-conjugated anti-Gr1 | R&D System | FAB1037F |
| Rat APC-conjugated anti-Ly6C | eBioscience | #175932 |
| Rat PE conjugated anti-Ly6G | BD Pharmingen | #551461 |
| Rat pacific blue conjugated anti-MHCII | BioLegend | #107620 |
| Rat eFluor660-conjugated anti-SOX2 | eBioscience | #50-9811-82 |
| Secondary |  |  |
| Anti-Mouse HRP | Jackson ImmunoResearch | 715-035-150 |
| Anti-Rabbit HRP | DakoCytomation | P0448 |
| Anti-Goat HRP | Jackson ImmunoResearch | 705-035-147 |
| Anti-Rat 488 | Thermo Fisher | A-212208 |
| Anti-Mouse 594 | Jackson ImmunoResearch | 715-585-151 |
| Anti-Goat 594 | Thermo Fisher | A-11058 |
| Anti-Mouse 488 | Jackson ImmunoResearch | 715 545 151 594 |
| Anti-Rabbit 594 | Thermo Fisher | R37119 |

**Table S2.**
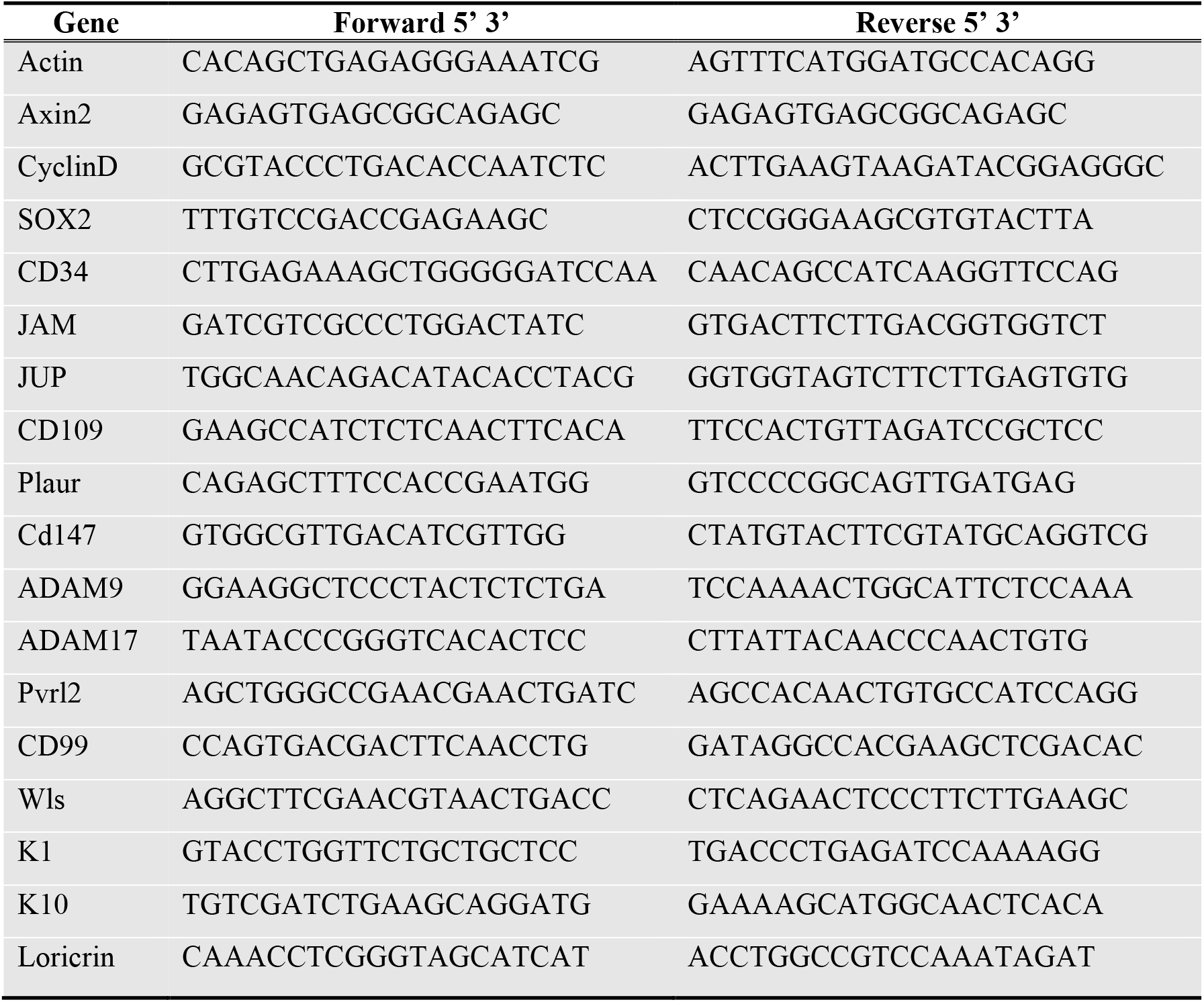
List of RT-qPCR primers.

## Supplementary Movie S1 legend

**Movie S1. Wnt-dependent cancer cell – macrophage attraction.**

Time-lapse microscopy of the wound area closure between HaCa4 mSCC cells (Tumor cells, TC) and RAW264.7 cells (Macrophages, MØ) under vehicle or IWP2 pre-treatment conditions. Recorded time 0 – 72 h. 15 frames per second.

## References

1 Sanchez-Danes, A. & Blanpain, C. Deciphering the cells of origin of squamous cell carcinomas. Nat Rev Cancer 18, 549–561, doi:10.1038/s41568-018-0024-5 (2018).

2 Nusse, R. & Clevers, H. Wnt/beta-Catenin Signaling, Disease, and Emerging Therapeutic Modalities. Cell 169, 985–999, doi:10.1016/j.cell.2017.05.016 (2017).

3 Malanchi, I. et al. Cutaneous cancer stem cell maintenance is dependent on beta-catenin signalling. Nature 452, 650–653, doi:10.1038/nature06835 (2008).

4 Zito, G. et al. Spontaneous tumour regression in keratoacanthomas is driven by Wnt/retinoic acid signalling cross-talk. Nat Commun 5, 3543, doi:10.1038/ncomms4543 (2014).

5 Schober, M. & Fuchs, E. Tumor-initiating stem cells of squamous cell carcinomas and their control by TGF-beta and integrin/focal adhesion kinase (FAK) signaling. Proc Natl Acad Sci U S A 108, 10544–10549, doi:10.1073/pnas.1107807108 (2011).

6 Oshimori, N. & Fuchs, E. The harmonies played by TGF-beta in stem cell biology. Cell Stem Cell 11, 751–764, doi:10.1016/j.stem.2012.11.001 (2012).

7 Oshimori, N., Oristian, D. & Fuchs, E. TGF-beta promotes heterogeneity and drug resistance in squamous cell carcinoma. Cell 160, 963–976, doi:10.1016/j.cell.2015.01.043 (2015).

8 Junankar, S. R., Eichten, A., Kramer, A., de Visser, K. E. & Coussens, L. M. Analysis of immune cell infiltrates during squamous carcinoma development. J Investig Dermatol Symp Proc 11, 36–43, doi:10.1038/sj.jidsymp.5650024 (2006).

9 Medler, T. R. & Coussens, L. M. Duality of the immune response in cancer: lessons learned from skin. J Invest Dermatol 134, E23–28, doi:10.1038/skinbio.2014.5 (2014).

10 Bottomley, M. J., Thomson, J., Harwood, C. & Leigh, I. The Role of the Immune System in Cutaneous Squamous Cell Carcinoma. Int J Mol Sci 20, doi:10.3390/ijms20082009 (2019).

11 Brown, S. et al. Correction of aberrant growth preserves tissue homeostasis. Nature 548, 334–337, doi:10.1038/nature23304 (2017).

12 Weber, C. et al. Macrophage Infiltration and Alternative Activation during Wound Healing Promote MEK1-Induced Skin Carcinogenesis. Cancer Res 76, 805–817, doi:10.1158/0008-5472.CAN-14-3676 (2016).

13 Fujimura, T., Kambayashi, Y., Fujisawa, Y., Hidaka, T. & Aiba, S. Tumor-Associated Macrophages: Therapeutic Targets for Skin Cancer. Front Oncol 8, 3, doi:10.3389/fonc.2018.00003 (2018).

14 Malsin, E. S., Kim, S., Lam, A. P. & Gottardi, C. J. Macrophages as a Source and Recipient of Wnt Signals. Front Immunol 10, 1813, doi:10.3389/fimmu.2019.01813 (2019).

15 Ojalvo, L. S., Whittaker, C. A., Condeelis, J. S. & Pollard, J. W. Gene expression analysis of macrophages that facilitate tumor invasion supports a role for Wnt-signaling in mediating their activity in primary mammary tumors. J Immunol 184, 702–712, doi:10.4049/jimmunol.0902360 (2010).

16 Debebe, A. et al. Wnt/beta-catenin activation and macrophage induction during liver cancer development following steatosis. Oncogene 36, 6020–6029, doi:10.1038/onc.2017.207 (2017).

17 Linde, N. et al. Macrophages orchestrate breast cancer early dissemination and metastasis. Nat Commun 9, 21, doi:10.1038/s41467-017-02481-5 (2018).

18 Taniguchi S, E. A., Van Duzer A, Kumar S, Leitenberger JJ, Oshimori N. Tumor-initiating cells establish an IL-33-TGF-β niche signaling loop to promote cancer progression. Science 6501, eaay1813, doi:doi: 10.1126/science.aay1813. (2020).

19 Ishida, Y. et al. Pivotal Involvement of the CX3CL1-CX3CR1 Axis for the Recruitment of M2 Tumor-Associated Macrophages in Skin Carcinogenesis. J Invest Dermatol, doi:10.1016/j.jid.2020.02.023 (2020).

20 Antsiferova, M. et al. Activin promotes skin carcinogenesis by attraction and reprogramming of macrophages. EMBO Mol Med 9, 27–45, doi:10.15252/emmm.201606493 (2017).

21 Linde, N. et al. Vascular endothelial growth factor-induced skin carcinogenesis depends on recruitment and alternative activation of macrophages. J Pathol 227, 17–28, doi:10.1002/path.3989 (2012).

22 Pettersen, J. S. et al. Tumor-associated macrophages in the cutaneous SCC microenvironment are heterogeneously activated. J Invest Dermatol 131, 1322–1330, doi:10.103/jid.2011.9 (2011).

23 Moussai, D. et al. The human cutaneous squamous cell carcinoma microenvironment is characterized by increased lymphatic density and enhanced expression of macrophage-derived VEGF-C. J Invest Dermatol 131, 229–236, doi:10.1038/jid.2010.266 (2011).

24 Boumahdi, S. et al. SOX2 controls tumour initiation and cancer stem-cell functions in squamouscell carcinoma. Nature 511, 246–250, doi:10.1038/nature13305 (2014).

25 Siegle, J. M. et al. SOX2 is a cancer-specific regulator of tumour initiating potential in cutaneous squamous cell carcinoma. Nat Commun 5, 4511, doi:10.1038/ncomms5511 (2014).

26 Holness, C. L. & Simmons, D. L. Molecular cloning of CD68, a human macrophage marker related to lysosomal glycoproteins. Blood 81, 1607–1613 (1993).

27 Abel, E. L., Angel, J. M., Kiguchi, K. & DiGiovanni, J. Multi-stage chemical carcinogenesis in mouse skin: fundamentals and applications. Nat Protoc 4, 1350–1362, doi:10.1038/nprot.2009.120 (2009).

28 Nassar, D., Latil, M., Boeckx, B., Lambrechts, D. & Blanpain, C. Genomic landscape of carcinogen-induced and genetically induced mouse skin squamous cell carcinoma. Nat Med 21, 946–954, doi:10.1038/nm.3878nm.3878 [pii] (2015).

29 DiGiovanni, J. Multistage carcinogenesis in mouse skin. Pharmacol Ther 54, 63–128, doi:10.1016/0163-7258(92)90051-z (1992).

30 Hausmann, G., Banziger, C. & Basler, K. Helping Wingless take flight: how WNT proteins are secreted. Nat Rev Mol Cell Biol 8, 331–336, doi:10.1038/nrm2141 (2007).

31 Carpenter, A. C., Rao, S., Wells, J. M., Campbell, K. & Lang, R. A. Generation of mice with a conditional null allele for Wntless. Genesis 48, 554–558, doi:10.1002/dvg.20651 (2010).

32 Li, S. et al. A keratin 15 containing stem cell population from the hair follicle contributes to squamous papilloma development in the mouse. Mol Carcinog 52, 751–759, doi:10.1002/mc.21896 (2013).

33 White, A. C. et al. Defining the origins of Ras/p53-mediated squamous cell carcinoma. Proc Natl Acad Sci USA 108, 7425–7430, doi:10.1073/pnas.1012670108 (2011).

34 Chae, W. J. & Bothwell, A. L. M. Canonical and Non-Canonical Wnt Signaling in Immune Cells. Trends Immunol 39, 830–847, doi:10.1016/j.it.2018.08.006 (2018).

35 Lien, W. H. & Fuchs, E. Wnt some lose some: transcriptional governance of stem cells by Wnt/beta-catenin signaling. Genes Dev 28, 1517–1532, doi:10.1101/gad.244772.114 (2014).

36 Farin, H. F. et al. Visualization of a short-range Wnt gradient in the intestinal stem-cell niche. Nature 530, 340–343, doi:10.1038/nature16937 (2016).

37 Cosin-Roger, J., Ortiz-Masia, M. D. & Barrachina, M. D. Macrophages as an Emerging Source of Wnt Ligands: Relevance in Mucosal Integrity. Front Immunol 10, 2297, doi:10.3389/fimmu.2019.02297 (2019).

38 Krimpenfort, P. et al. A natural WNT signaling variant potently synergizes with Cdkn2ab loss in skin carcinogenesis. Nat Commun 10, 1425, doi:10.1038/s41467-019-09321-8 (2019).

39 Long, A. et al. WNT10A promotes an invasive and self-renewing phenotype in esophageal squamous cell carcinoma. Carcinogenesis 36, 598–606, doi:10.1093/carcin/bgv025 (2015).

40 Caulin, C. et al. Suppression of the metastatic phenotype of a mouse skin carcinoma cell line independent of E-cadherin expression and correlated with reduced Ha-ras oncogene products. Mol Carcinog 15, 104–114, doi:10.1002/(SICI)1098-2744(199602)15:2<104::AID-MC3>3.0.CO;2-J (1996).

41 Raschke, W. C., Baird, S., Ralph, P. & Nakoinz, I. Functional macrophage cell lines transformed by Abelson leukemia virus. Cell 15, 261–267, doi:10.1016/0092-8674(78)90101-0 (1978).

42 Chen, B. et al. Small molecule-mediated disruption of Wnt-dependent signaling in tissue regeneration and cancer. Nat Chem Biol 5, 100–107, doi:10.1038/nchembio.137 (2009).

43 Sunagawa, M. et al. Suppression of skin tumorigenesis in CD109-deficient mice. Oncotarget 7, 82836–82850, doi:10.18632/oncotarget.12653 (2016).

44 Samanta, D. et al. Structure of Nectin-2 reveals determinants of homophilic and heterophilic interactions that control cell-cell adhesion. Proc Natl Acad Sci USA 109, 14836–14840, doi:10.1073/pnas.1212912109 (2012).

45 Pasello, M., Manara, M. C. & Scotlandi, K. CD99 at the crossroads of physiology and pathology. J Cell Commun Signal 12, 55–68, doi:10.1007/s12079-017-0445-z (2018).

46 Manara, M. C., Pasello, M. & Scotlandi, K. CD99: A Cell Surface Protein with an Oncojanus Role in Tumors. Genes (Basel) 9, doi:10.3390/genes9030159 (2018).

47 Zucchini, C. et al. CD99 suppresses osteosarcoma cell migration through inhibition of ROCK2 activity. Oncogene 33, 1912–1921, doi:10.1038/onc.2013.152 (2014).

48 Mannion, A. J., Odell, A.F, Taylor, A, Jones, P.J, Cook, G.P. CD99 regulates cancer cell transendothelial migration and endothelial cell function via CDC42 and actin remodelling. bioRxiv 760934, doi:doi: https://doi.org/10.1101/760934 (2019).

49 Schenkel, A. R., Mamdouh, Z., Chen, X., Liebman, R. M. & Muller, W. A. CD99 plays a major role in the migration of monocytes through endothelial junctions. Nat Immunol 3, 143–150, doi:10.1038/ni749 (2002).

50 Goswami, D. et al. Endothelial CD99 supports arrest of mouse neutrophils in venules and binds to neutrophil PILRs. Blood 129, 1811–1822, doi:10.1182/blood-2016-08-733394 (2017).

51 Li, Y. T. et al. Blood flow guides sequential support of neutrophil arrest and diapedesis by PILR-beta1 and PILR-alpha. Elife 8, doi:10.7554/eLife.47642 (2019).

52 Choi, G., Roh, J. & Park, C. S. CD99 Is Strongly Expressed in Basal Cells of the Normal Adult Epidermis and Some Subpopulations of Appendages: Comparison with Developing Fetal Skin. J Pathol Transl Med 50, 361–368, doi:10.4132/jptm.2016.06.19 (2016).

53 Beck, B. et al. A vascular niche and a VEGF-Nrp1 loop regulate the initiation and stemness of skin tumours. Nature 478, 399–403, doi:10.1038/nature10525 (2011).

54 Miao, Y. et al. Adaptive Immune Resistance Emerges from Tumor-Initiating Stem Cells. Cell 177, 1172–1186 e1114, doi:10.1016/j.cell.2019.03.025 (2019).

55 Wellenstein, M. D. & de Visser, K. E. Cancer-Cell-Intrinsic Mechanisms Shaping the Tumor Immune Landscape. Immunity 48, 399–416, doi:10.1016/j.immuni.2018.03.004 (2018).

56 Sainz, B., Jr., Carron, E., Vallespinos, M. & Machado, H. L. Cancer Stem Cells and Macrophages: Implications in Tumor Biology and Therapeutic Strategies. Mediators Inflamm 2016, 9012369, doi:10.1155/2016/9012369 (2016).

57 Lu, H. et al. A breast cancer stem cell niche supported by juxtacrine signalling from monocytes and macrophages. Nat Cell Biol 16, 1105–1117, doi:10.1038/ncb3041 (2014).

## Methods References

1 Wang, F. et al. RNAscope: a novel in situ RNA analysis platform for formalin-fixed, paraffin-embedded tissues. J Mol Diagn 14, 22–29, doi:10.1016/j.jmoldx.2011.08.002 S1525-1578(11)00257-1 [pii] (2012).

2 Holness, C. L. & Simmons, D. L. Molecular cloning of CD68, a human macrophage marker related to lysosomal glycoproteins. Blood 81, 1607–1613 (1993).

3 Boumahdi, S. et al. SOX2 controls tumour initiation and cancer stem-cell functions in squamous-cell carcinoma. Nature 511, 246–250, doi:10.1038/nature13305 (2014).

4 Siegle, J. M. et al. SOX2 is a cancer-specific regulator of tumour initiating potential in cutaneous squamous cell carcinoma. Nat Commun 5, 4511, doi:10.1038/ncomms5511 (2014).

5 Morris, R. J. et al. Capturing and profiling adult hair follicle stem cells. Nature biotechnology 22, 411–417, doi:10.1038/nbt950 (2004).

6 Clausen, B. E., Burkhardt, C., Reith, W., Renkawitz, R. & Forster, I. Conditional gene targeting in macrophages and granulocytes using LysMcre mice. Transgenic research 8, 265–277 (1999).

7 Carpenter, A. C., Rao, S., Wells, J. M., Campbell, K. & Lang, R. A. Generation of mice with a conditional null allele for Wntless. Genesis 48, 554–558, doi:10.1002/dvg.20651 (2010).

8 Ferrer-Vaquer, A. et al. A sensitive and bright single-cell resolution live imaging reporter of Wnt/ss-catenin signaling in the mouse. BMC Dev Biol 10, 121, doi:10.1186/1471-213X-10-121 1471-213X-10-121 [pii] (2010).

9 Abel, E. L., Angel, J. M., Kiguchi, K. & DiGiovanni, J. Multi-stage chemical carcinogenesis in mouse skin: fundamentals and applications. Nat Protoc 4, 1350–1362, doi:10.1038/nprot.2009.120 (2009).

10 Nassar, D., Latil, M., Boeckx, B., Lambrechts, D. & Blanpain, C. Genomic landscape of carcinogen-induced and genetically induced mouse skin squamous cell carcinoma. Nat Med 21, 946–954, doi:10.1038/nm.3878 nm.3878 [pii] (2015).

11 White, A. C. et al. Defining the origins of Ras/p53-mediated squamous cell carcinoma. Proc Natl Acad Sci U S A 108, 7425–7430, doi:10.1073/pnas.1012670108 (2011).

12 Babij, P. et al. “Blue heart”: characterization of a mifepristone-dependent system for conditional gene expression in genetically modified animals. Biochim Biophys Acta 1627, 15–25, doi:S0167478103000526 [pii] (2003).

13 Raschke, W. C., Baird, S., Ralph, P. & Nakoinz, I. Functional macrophage cell lines transformed by Abelson leukemia virus. Cell 15, 261–267, doi:10.1016/0092-8674(78)90101-0 (1978).

14 Caulin, C. et al. Suppression of the metastatic phenotype of a mouse skin carcinoma cell line independent of E-cadherin expression and correlated with reduced Ha-ras oncogene products. Mol Carcinog 15, 104–114, doi:10.1002/(SICI)1098-2744(199602)15:2<104::AID-MC3>3.0.CO;2-J (1996).

15 Llorens, A. et al. Down-regulation of E-cadherin in mouse skin carcinoma cells enhances a migratory and invasive phenotype linked to matrix metalloproteinase-9 gelatinase expression. Lab Invest 78, 1131–1142 (1998).

16 Shahbazi, M. N. et al. CLASP2 interacts with p120-catenin and governs microtubule dynamics at adherens junctions. J Cell Biol 203, 1043–1061, doi:10.1083/jcb.201306019 (2013).

17 Fontenete, S. & Perez-Moreno, M. Isolation of Cancer Stem Cells from Squamous Cell Carcinoma. Methods Mol Biol 1879, 407–414, doi:10.1007/7651_2018_162 (2019).

18 Donoghue, P. M., Hughes, C., Vissers, J. P., Langridge, J. I. & Dunn, M. J. Nonionic detergent phase extraction for the proteomic analysis of heart membrane proteins using label-free LC-MS. Proteomics 8, 3895–3905, doi:10.1002/pmic.200800116 (2008).

19 Wieczorek, S. et al. DAPAR & ProStaR: software to perform statistical analyses in quantitative discovery proteomics. Bioinformatics 33, 135–136, doi:10.1093/bioinformatics/btw580 (2017).

20 Bo, T. H., Dysvik, B. & Jonassen, I. LSimpute: accurate estimation of missing values in microarray data with least squares methods. Nucleic acids research 32, e34, doi:10.1093/nar/gnh026 (2004).

